# Th17 effector cytokines induce shared and distinct microglial and endothelial cell responses in post-streptococcal encephalitis

**DOI:** 10.64898/2026.02.04.703836

**Authors:** Charlotte R. Wayne, Uğur Akcan, Travis E. Faust, Violeta Durán-Laforet, Danny Jamoul, Luca Bremner, Nicole Ampatey, Büşra T. Akcan, Sarah J. Ho, Bogoljub Ciric, Shannon L. Delaney, Wendy S. Vargas, Susan Swedo, Vilas Menon, Dorothy P. Schafer, Tyler Cutforth, Dritan Agalliu

## Abstract

Group A *Streptococcus* (GAS) infections can lead to neuropsychiatric sequelae in children, yet the mechanisms driving post-infectious brain pathology remain poorly defined. In a mouse disease model, Th17 lymphocytes induce microglial activation, blood–brain barrier (BBB) dysfunction, and neural circuit impairment; however, the transcriptional programs underlying these effects, and the specific Th17-derived cytokines involved are unclear. Using mouse genetics, single-cell RNA sequencing, and spatial transcriptomics, we show that GAS infections induce inflammatory gene programs in microglia and brain endothelial cells (BECs), accompanied by downregulation of BBB-associated transcripts in BECs. Spatial transcriptomic analyses reveal that GAS-responsive microglia are enriched near infiltrating T cells. Several chemokines upregulated in microglia following GAS infection in mice are elevated in sera from affected patients. Conditional ablation of GM-CSF in CD4^+^ T cells partially attenuates microglial chemokine gene expression, but does not restore BBB integrity. Neutralization of IL-17A partially rescues BBB transcriptional changes in BECs and reduces microglial chemokine expression; however, compensatory peripheral immune responses associated with persistent infection exacerbate BBB disruption. In contrast, microglia/macrophage-specific deletion of IL-17 receptor A partially rescues BBB deficits following GAS infection. Together, these findings identify IL-17A–IL-17RA signaling in microglia as a critical driver of BBB dysfunction after GAS infections.

## Main

Neuropsychiatric and cognitive disturbances can arise from bacterial or viral infections, even in the absence of direct brain infection [1]. Peripheral infections with *Streptococcus pyogenes*, or Group A *Streptococcus* (GAS), can give rise to secondary sequelae that impair central nervous system (CNS) function, including movement disorders such as Sydenham’s chorea (SC), and psychiatric syndromes such as Pediatric Autoimmune Neuropsychiatric Disorders Associated with Streptococcal infections (PANDAS; reviewed in [2, 3]). SC is characterized by involuntary, uncoordinated movements and behavioral abnormalities, whereas PANDAS presents with a spectrum of psychiatric and fine motor symptoms, including obsessive–compulsive behaviors, vocal or motor tics, reduced appetite or anorexia nervosa, and severe separation anxiety [2, 3]. The mechanisms driving CNS sequelae following GAS infections remain poorly understood, but are thought to involve aberrant anti-pathogen immune responses that target the CNS (reviewed in [3–5]), leading to neuroinflammation, blood-brain barrier (BBB) disruption, and neuronal circuitry dysfunction (reviewed in [5, 6]).

Neuroimaging studies have identified increased basal ganglia volume in both SC and PANDAS patients [7, 8], and increased microglial/astrocytic activation in the basal ganglia of PANDAS patients compared to healthy controls [9]. Autoantibodies targeting basal ganglia structures have also been described in both disorders, including antibodies against dopamine D2 receptors in SC [10, 11], anti-dopamine 1 receptor (D1R) antibodies in PANDAS [12], and antibodies against striatal cholinergic interneurons and other neuronal targets in PANDAS [13–16]. Nevertheless, the molecular mechanisms underlying this brain pathology—here referred to as post-streptococcal basal ganglia encephalitis (post-GAS-BGE)—remain incompletely defined. Elucidating these mechanisms is critical for improving diagnosis, which has been hampered by the lack of reliable biomarkers [3, 17–19], and for developing effective treatments for the chronic phase of these disorders [20, 21].

T helper 17 (Th17) cells producing interleukin-17A (IL-17A) play an essential role in host defense against extracellular pathogens, and repeated intranasal GAS infections elicit robust Th17 responses in both mice and humans [22–24]. Using a juvenile mouse model of intranasal GAS infection, we previously showed that CD4^⁺^ T cells, including Th17 and Th1 subsets, migrate from the nasal cavity into the anterior brain, where they localize predominantly to the olfactory bulb (OB), the brain’s primary relay for olfactory input. This process is accompanied by BBB disruption, microglial activation, and degradation of excitatory synapses, leading to anosmia and abnormal odor-evoked responses [23, 25]. Genetic elimination of Th17 cells partially rescues BBB dysfunction, microglial activation, and olfactory circuit abnormalities in this mouse model [25], indicating a central role for Th17 lymphocytes in post-GAS-BGE pathology. However, the transcriptional responses of microglia, brain endothelial cells (BECs), and other CNS cell types following intranasal GAS infection remain unknown.

Another key unresolved question is how Th17 effector cytokines influence inflammatory responses in microglia and BECs after GAS infections. Th17 cells exhibit considerable phenotypic plasticity [26, 27], and chronic inflammation can drive a transition from conventional IL-17A-producing Th17 cells to a population that produces interferon-γ (IFNγ) and granulocyte-macrophage colony-stimulating factor (GM-CSF), and expresses the transcription factor T-bet [28]. These pathogenic Th17 cells (Th17^path^) are required for disease development in the experimental autoimmune encephalomyelitis (EAE), a mouse model for multiple sclerosis (MS) [29, 30], and are associated with disease severity in several human autoimmune disorders [31–33]. Despite the established role of GM-CSF in pathogenic Th17-mediated tissue damage, it remains unclear whether GM-CSF is produced by CD4^⁺^ T cells infiltrating the brain after GAS infections, and how GM-CSF and IL-17A differentially contribute to transcriptional changes in microglia and BECs and to CNS pathology in this rodent model.

To address these questions, we combined mouse genetic approaches with single-cell RNA sequencing, spatial transcriptomics, and targeted validation studies in a mouse model of SC/PANDAS. We identify extensive transcriptional changes in both microglia and BECs following repeated GAS infection, including downregulation of microglial homeostatic genes and BBB-associated genes in BECs, alongside upregulation of interferon-response, chemokine, and antigen-presentation programs in both cell types. Spatial transcriptomic analyses reveal enrichment of Streptococcus-responsive microglia within the glomerular layer of the OB, in close proximity to infiltrating CD4^⁺^T cells. Conditional ablation of GM-CSF in CD4^⁺^ T cells partially attenuates microglial chemokine gene expression, but does not restore BBB integrity. Systemic neutralization of IL-17A partially rescues BBB-associated transcriptional changes in BECs and reduces microglial chemokine expression; however, compensatory peripheral immune responses associated with persistent GAS infection exacerbate BBB disruption. In contrast, microglia/macrophage-specific deletion of IL-17 receptor A partially ameliorates BBB deficits following GAS infection. Together, these findings support a role for IL-17A–IL-17RA signaling in microglia and macrophages in shaping BBB dysfunction after GAS infection, and suggest that targeting this pathway may complement existing therapeutic strategies for chronic SC/PANDAS in the absence of active infection [20, 21].

## Results

### Microglia and BECs show major transcriptional shifts after multiple GAS infections

We have shown that multiple intranasal GAS infections induce infiltration of CD4^+^ T cells in the OB, accompanied by microglial activation, BBB damage and degradation of excitatory synapses, leading to aberrant odor-evoked neural circuit responses [23, 25]. To investigate, at the transcriptional level, how CNS cell types respond to GAS infections, we isolated and profiled OB cells from P60 mice 18 hours after the fifth GAS infection using scRNAseq (**Figure 1a; Supplementary Data Table 1**). Mice inoculated with PBS served as controls.

**Figure 1.**
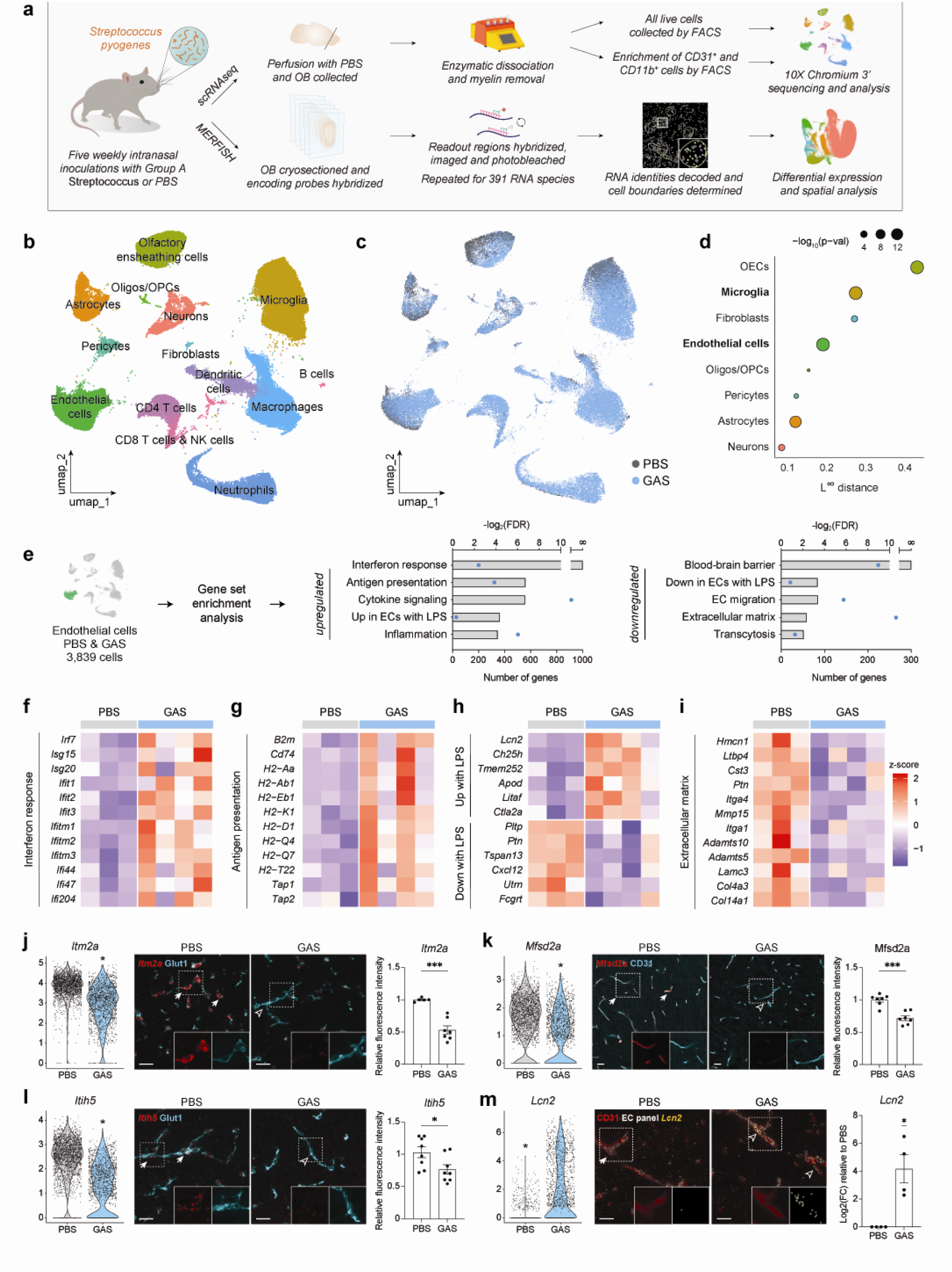
Brain endothelial cells upregulate inflammatory signatures and downregulate BBB marker expression following GAS infections. **a**, Experimental workflow for single-cell RNA sequencing (scRNAseq) and MERFISH experiments. **b-c**, UMAP visualization of olfactory bulb (OB) scRNAseq from PBS controls (gray) and GAS-infected mice (blue), with enrichment of CD31^+^ brain endothelial cells (BECs) and CD11b^+^ (myeloid cells). **d**, Quantification of transcriptional shifts across OB cell types between PBS and GAS conditions using L^∞^ distance (cross-entropy test). **e**, Gene set enrichment analysis (GSEA) of differentially expressed genes in BECs after GAS infection. Inflammatory and interferon-related pathways are enriched, whereas blood-brain barrier (BBB) gene sets are depleted. Bars show significance of enrichment (-log_2_FDR) and blue dots indicate the number of significant genes per set. **f-i**, Heatmaps of interferon response, antigen presentation, endothelial LPS response and extracellular matrix in BECs from PBS and GAS conditions [log(z-score)]. The red colors mean a high z-score and the blue colors mean a low z-score. Significant genes are in balck (adjusted p < 0.05) and non-significant ones are in gray (adjusted p > 0.05). **j-m**, Downregulation of BBB-associated transcripts (*Itm2a*, *Itih5* and *Mfsd2a*) in BECs measured by scRNAseq (left; Wilcoxon Rank Sum test, *adjusted p < 0.05). Representative images (center) and quantification (right) of changes in the BBB transcripts by fluorescence *in situ* RNA hybridization (FISH) for *Itm2a*, *Itih5* mRNAs (red) combined with immunofluorescence for BEC marker Glut1 (**j-l**; cyan) and immunofluorescence staining for Mfsd2a (red) and BEC marker CD31 (**m**; red). Scale bars = 25 μm. Comparisons were performed with unpaired t test with Welch’s correction (* p < 0.05; n = 7-9 per condition). Data are mean +/- SEM.

Following quality filtering and batch correction, OB cells were visualized using Uniform Manifold Approximation and Projection (UMAP), and cluster identities were assigned based on established cell-type markers (**Figure 1b, c; Extended Data Figure 1a**). Cluster annotations were further refined, and were consistent across biological replicates (**Extended Data Figure 1e**). A cross entropy test [34] comparing PBS and GAS conditions revealed the largest shifts in olfactory ensheathing cells (OECs), consistent with their capacity to recruit immune cells during intranasal inflammation (**Figure 1d, Supplementary Table 2**) [35]. Microglia and BECs also exhibited significant transcriptome shifts **(Figure 1d, Supplementary Table 2)**. In contrast, astrocytes, - key mediators of inflammation in many neuroinflammatory models [36], - displayed only modest transcriptome changes after GAS infections **(Figure 1d, Supplementary data Table 2)**. Given the prominent BBB dysfunction and microglial activation observed in this disease model [23, 25], we focused subsequent analyses on BECs and microglia. To increase their representation in scRNAseq datasets, CD31^+^ BECs and CD11b^+^ myeloid cells were enriched by fluorescence-activated cell sorting (FACS) prior to sequencing **(Extended Data Figure 1b–d)**.

### BECs upregulate inflammatory programs and downregulate BBB-related transcripts after GAS infections

To define BEC-specific transcriptional responses to GAS infections, BECs were extracted from the scRNAseq dataset and subclustered (**Extended Data Figure 1f, g**). Differentially expressed genes (DEGs) were analyzed using gene set enrichment analysis (GSEA) [37, 38] with curated gene lists (**Supplementary Data Tables 3, 4**). Compared to PBS controls, BECs from GAS-infected mice showed significant upregulation of genes associated with interferon response, antigen presentation, cytokine signaling, inflammation and endothelial cell (ECs) response to lipopolysaccharide (LPS) [39], alongside downregulation of BBB-associated genes [40] (**Figure 1e-h; Extended Data Figure 1h-k; Supplementary Data Table 3**). Within the BBB transcriptome, transcripts related to adherens and tight junction, transporters, regulators of transcytosis and brain endothelial identity were significantly reduced in the GAS condition (**Extended Data Figure 1h-k**). Decreased expression of two BBB-specific genes, *Itm2a* and *Itih5* [40] as well as increased expression of *lipocalin 2 (Lcn2)*, an IL-17A-induced inflammatory mediator [41, 42], was validated by fluorescence *in situ* hybridization (FISH) and MERFISH (see below), respectively, in OB sections from GAS-infected mice (**Figure 1j, l, m**). Consistent with increased serum IgG transport across the compromised BBB in GAS-infected mice [23, 25], the transcytosis suppressor gene *Mfsd2a* [43] was downregulated at both the mRNA and protein levels in BECs (**Figure 1k; Extended Data Figure 1k**). Unexpectedly, genes promoting caveolar transport (including *Cav1-2, Cavin1-3*) were also downregulated in BECs from GAS-infected mice (**Extended Data Figure 1j**). Mixed-effects modeling analysis of BEC DEGs revealed also significant downregulation of several extracellular matrix (ECM) genes (**Figure 1i**), critical for BBB integrity (reviewed in [44]).

### Microglia upregulate inflammatory and antigen-presentation programs after GAS infections

GSEA of microglial DEGs revealed enrichment of disease-associated microglia (DAM) signatures, antigen-presentation pathways, cytokine and growth factor signaling, and interferon response genes in GAS compared to PBS conditions (**Figure 2a-c; Extended Data Figure 2a; Supplementary Data Table 3**). Consistent with these findings, flow cytometry analysis showed increased expression of the antigen-presentation protein CD74, as well as TNF and CCL5 cytokines in microglia from GAS-infected mice, accompanied by reduced expression of homeostatic receptors CX3CR1 and P2RY12 (**Figure 2d-i**). Elevated levels of CCL2, CCL4, CCL5, CXCL10 and TNF proteins were also detected in whole OB lysates from GAS-infected mice using a multiplex immunoassay (**Extended Data Figure 2c**). Importantly, upregulation of *Ccl3* and *Ccl4* transcripts in microglia was not attributable to enzymatic dissociation artifacts [45] as their expression did not correlate with an *ex vivo* activation gene signature (**Extended Data Figure 2d, e**).

**Figure 2.**
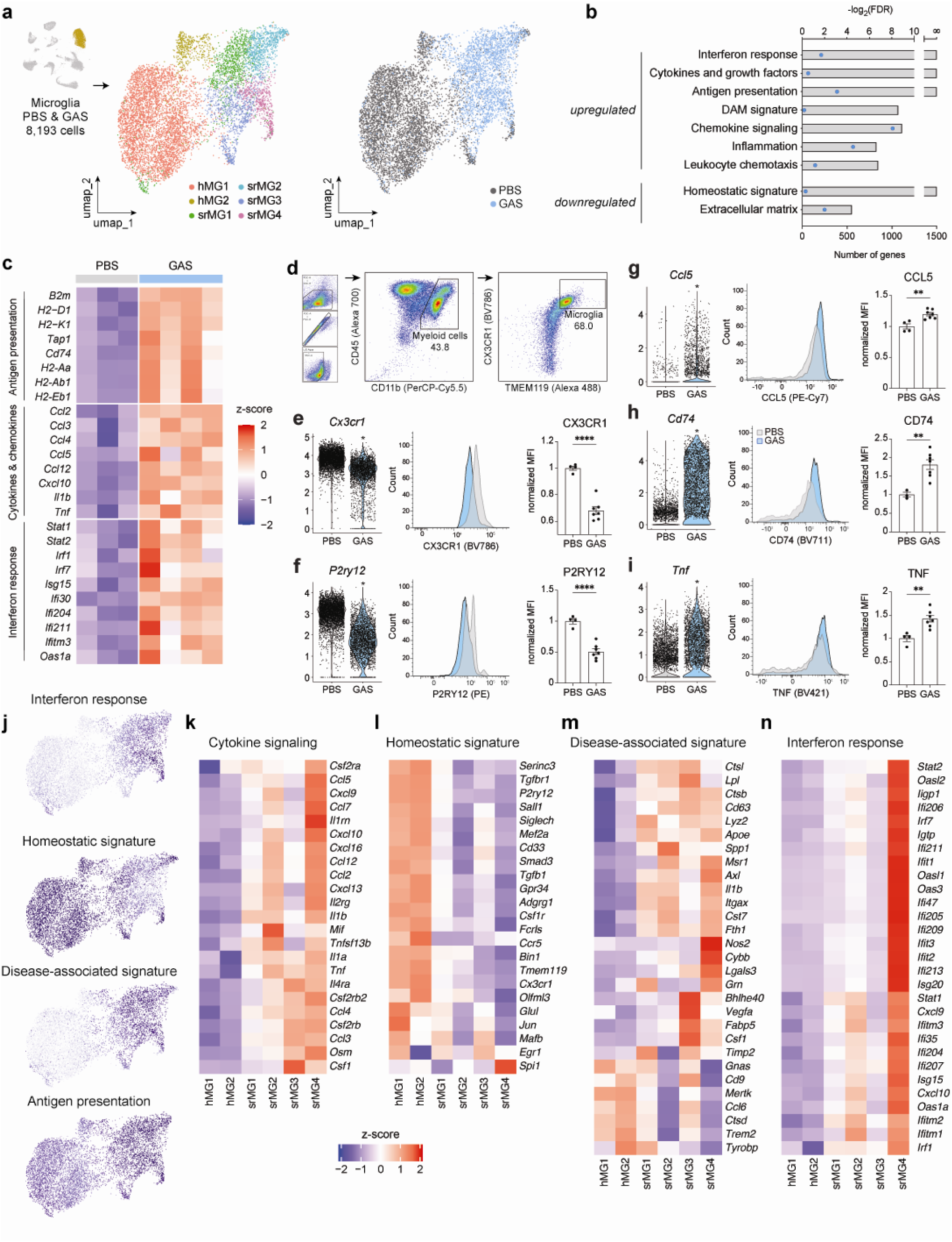
Microglia upregulate inflammatory and disease-associated transcriptional programs following recurrent GAS infections. **a,** UMAP visualization of microglia from olfactory bulbs (OBs) of PBS- and GAS-treated mice identifies six transcriptional clusters: two homeostatic populations (hMG1–2) and four Streptococcus-responsive subtypes (srMG1–4). Cells are colored by cluster (left) or condition (right; PBS, gray; GAS, blue). **b,** Gene set enrichment analysis (GSEA) of differentially expressed genes in microglia after GAS infection. Bars represent –log₂(FDR); dots indicate the number of significant genes per pathway. **c,** Heatmap of antigen presentation, cytokine/chemokine, and interferon-response genes in PBS versus GAS microglia. **d–i,** Differential expression of activation markers (*Cd74, Tnf, Ccl5*) and homeostatic receptors (*Cx3cr1, P2ry12*) measured by scRNAseq (left; Wilcoxon rank-sum test, adjusted *p* < 0.05) and validated by flow cytometry (center; representative histograms; right, quantification of normalized median fluorescence intensity). Statistical comparisons were performed using unpaired *t* tests (\**p* < 0.01; \*\*\**p* < 0.0001). **j,** Feature plots showing expression of selected homeostatic, disease-associated, antigen-presentation, and interferon-response genes across microglial populations. **k–n,** Heatmaps of cytokine signaling, homeostatic, disease-associated microglia (DAM), and interferon-response gene modules across the six microglial clusters (log z-score). Significant genes are labeled in black (adjusted *p* < 0.05).

To further resolve microglial heterogeneity, scRNAseq data from PBS and GAS conditions were subclustered revealing six distinct microglial populations (**Supplementary Data Table 3**). Two clusters (hMG1-2), comprised predominantly of PBS-derived cells, expressed canonical homeostatic microglial genes (**Figure 2j, l**). In contrast, four *Streptococcus*-responsive clusters (srMG1-4) exhibited graded upregulation of cytokine signaling, DAM-associated genes, and antigen-presentation programs (**Figure 2j, k, m, n**). For example, while all srMG clusters expressed *Ccl3* and *Ccl4*, *Ccl2*, *Ccl5*, and *Tnf* were most highly expressed in srMG2 and srMG4 (**Figure 2k**). The srMG4 cluster showed the strongest enrichment of interferon-response genes (**Figure 2n**).

Finally, the scRNAseq analysis revealed an increased abundance of macrophages in OBs from GAS-infected mice (**Figure 1b, c, Extended Data Figure 1c, d**). To determine whether peripheral macrophages infiltrate the brain parenchyma, *CX3CR1^GFP^* transgenic mice [46] were crossed to a *TMEM119^tdTomato^*reporter mice [47]. This strategy distinguishes resident microglia (GFP^+^ tdTomato^+^) from peripheral macrophages (GFP^+^ tdTomato^−^). Macrophages were confined to perivascular and meningeal regions in both PBS- and GAS-infected mice OBs, with minimal parenchymal infiltration (**Extended Data Figure 2f-j**), although both populations were increased in GAS-infected OBs (**Extended Data Figure 2i, j**). Thus, peripheral macrophages accumulate near vascular and meningeal compartments, but do not invade the brain parenchyma following repeated GAS infections **(Extended Data Figure 2k)**.

### *Streptococcus*-responsive microglia are enriched in the glomerular OB layer in close proximity to T cells

To understand the regional distribution of “*Streptococcus*-responsive” microglial clusters and their relationship to infiltrating T cells, we performed spatial transcriptomics using multiplexed error-robust fluorescence in situ hybridization (MERFISH) [48] on OB sections (**Figure 3a, b**). We probed 391 genes primarily expressed by microglia and BECs (See Methods), as these cell types showed robust transcriptional shifts by scRNAseq **(Figure 1d)**. Cell-type identities were assigned using both canonical markers and spatial localization **(Extended Data Figure 3a-c)**. MERFISH identified cell types consistent with those detected by scRNAseq **(Figure 3a-c)**, while providing improved representation of neuronal populations compared to FACS-based scRNAseq **(Figure 3a, b; Extended Data Figure 1c, 3a-c)**.

**Figure 3.**
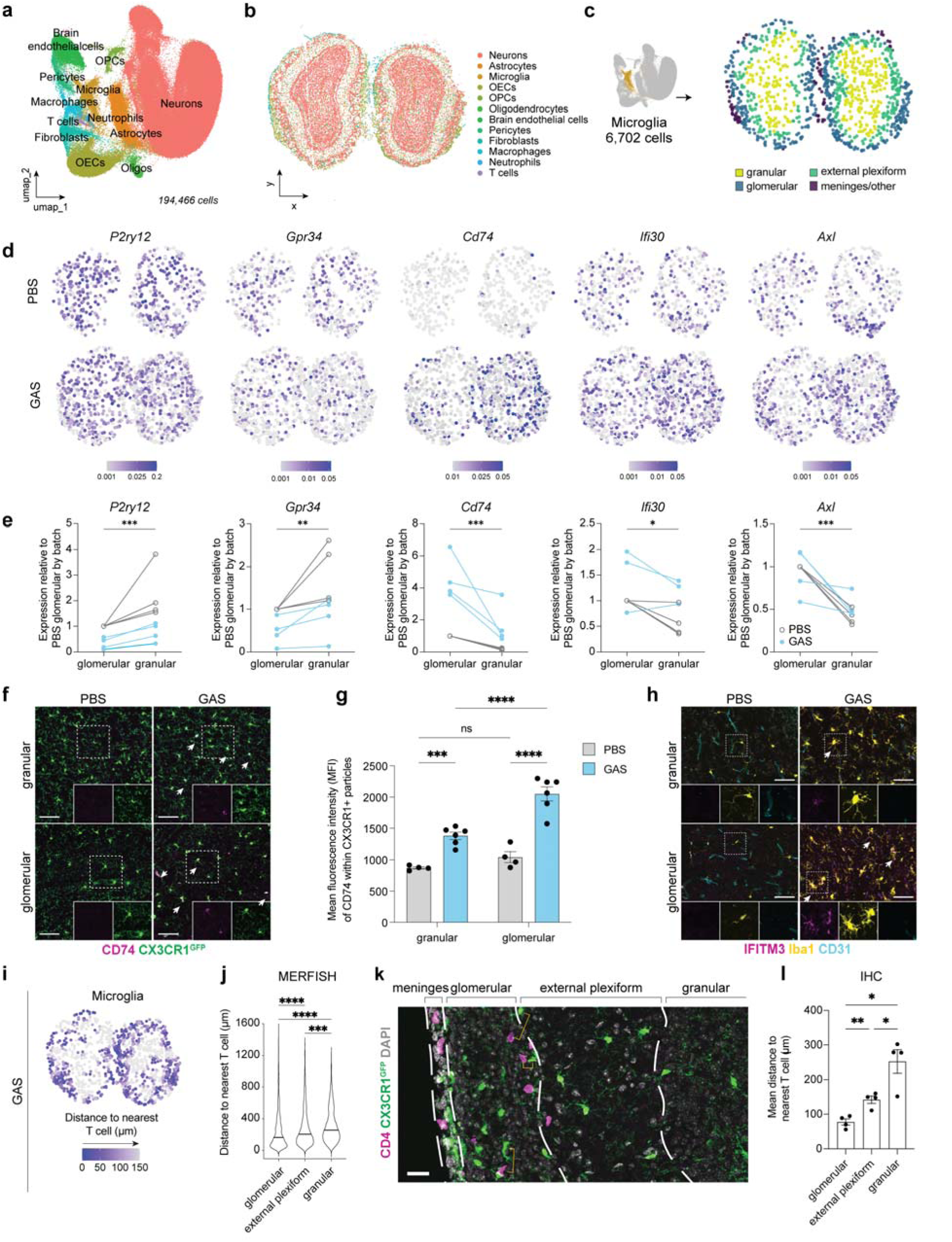
Spatial transcriptomics reveals enhanced Streptococcus-responsive microglial activation near infiltrating T cells. **a, b,** UMAP visualization (**a**) and representative spatial coordinate plot (**b**) of all MERFISH samples. **c,** Spatial map of microglia annotated by olfactory bulb (OB) layer. **d,** Representative MERFISH expression maps for homeostatic (*P2ry12, Gpr34*) and Streptococcus-responsive (*Cd74, Ifi30, Axl*) transcripts in PBS and GAS OBs (n = 4 mice per group). **e,** Relative expression of homeostatic and GAS-responsive genes in microglia from the glomerular versus granular OB layers. Values were normalized to PBS glomerular expression within batch. Statistical comparisons were performed using paired ratio *t* tests (*p* < 0.05; \**p* < 0.01; \*\*\**p* < 0.0001; n = 4 mice per group). **f, g,** Representative immunofluorescence (IF) images of CD74⁺ myeloid cells (red; arrows) in *CX3CR1^eGFP+^*mice (**f**) and quantification across OB layers (**g**). Data are shown as mean ± SEM. Statistical analysis was performed using two-way ANOVA with Šídák’s multiple comparisons (ns, p > 0.05; \**p* < 0.01; \*\**p* < 0.001; \*\*\**p* < 0.0001; n = 4–6 mice per group). **h,** Representative IF images of IFITM3 (red), Iba1 (yellow; myeloid cells), and CD31 (blue; vessels) in glomerular and granular OB layers from PBS and GAS-infected mice. **i, j,** MERFISH-based spatial mapping (**i**) and quantification (**j**) of microglial distance to the nearest CD4⁺ T cell across OB layers in PBS and GAS conditions. **k, l,** Representative IF images of CD4⁺ T cells (red) and *CX3CR1^eGFP+^* microglia (**k**) and quantification of mean intercellular distances (**l**). Statistical comparisons were performed using one-way ANOVA (ns, p > 0.05; \**p* < 0.01; \*\**p* < 0.001; \*\*\**p* < 0.0001; n = 4 mice per group).

MERFISH revealed a distinct spatial pattern of Streptococcus-responsive gene expression across OB layers. Transcripts for *Cd74*, *Ifi30* and *Axl* (pathways upregulated in GAS microglia) were expressed at higher levels in microglia located in the glomerular compared to the granular layer of the OB (**Figure 3c-e**). Immunofluorescence (IF) analysis confirmed increased CD74 protein expression, critical for proper folding and trafficking of MHC class II molecules for antigen presentation (reviewed in [49]), in glomerular-layer microglia, mirroring the spatial transcriptomic pattern **(Figure 3f, g)**. IFITM3, a key interferon-response protein [50], similarly showed higher expression in glomerular microglia by IF (**Figure 3h**), consistent with *Ifi30* mRNA distribution by MERFISH. In contrast, homeostatic genes (*P2ry12*, *Gpr34*) were modestly enriched in granular-layer microglia **(Figure 3d, e)**.

Given that infiltrating CD4^⁺^ T cells accumulate predominantly in the glomerular layer and meninges [23, 25], we quantified microglia-T cell proximity. Distances between microglia and CD4^⁺^ T cells were significantly shorter in the glomerular and external plexiform layers compared to the granular layer, as assessed by MERFISH and IF (**Figure 3i-l**). Together, these data suggest that microglia in the glomerular layer undergo more pronounced transcriptional activation after GAS infections, correlating with their proximity to infiltrating CD4^⁺^ T cells.

### GAS infection alters predicted cell–cell communication networks in the OB

To assess whether GAS infection alters intercellular communication, we analyzed scRNAseq data using CellChat [51, 52]. CD4^⁺^ T cells were predicted to communicate primarily with microglia, olfactory ensheathing cells (OECs), and BECs, while microglia showed predicted interactions with macrophages and CD4^⁺^T cells (**Extended Data Figure 3e**). Inferred ligand - receptor interactions included both secreted and contact-dependent pathways. Microglia - BEC interactions were predicted to involve adhesion and inflammatory signaling molecules (e.g., *Jama*, *Tnf*, *Vegfb*), whereas CD4^⁺^ T cell - microglia communication was predicted to involve cytokines and chemokines (e.g., *Ifng*, *Tnf*, *Ccl5*) as well as adhesion molecules (**Extended Data Fig. 3f)**.

### PANDAS/PANS patients display elevated serum levels of inflammatory cytokines and growth factors

Previous studies have reported microglial and astrocytic activation in the basal ganglia of PANDAS patients [9] and elevated levels of select cytokines (e.g. IL-17A, TNF) in PANDAS/PANS patients sera [53, 54], but the breadth of systemic inflammatory changes during the acute disease phase remains unclear. Given the elevated cytokine and chemokine expression observed in the OB of GAS-infected mice (**Figure 2c–i; Extended Data Figure 2c**), we analyzed serum samples from PANDAS/PANS patients and controls. Serum was collected from 10 PANDAS patients enrolled in an intravenous immunoglobulin (IVIg) clinical trial at the National Institute for Mental Health (NIMH) [55], 13 PANDAS/PANS cases recruited at Columbia University Irving Medical Center (CUIMC) and 11 age-and sex-matched controls obtained from the NIMH (**Supplementary Data Table 7**). Using an unbiased multiplex immunoassay profiling 45 analytes (**Supplementary Data Table 6**), we identified significant elevations in 13 cytokines, chemokines and growth factors in patient sera (**Table 1**; adjusted *p* < 0.05). Six cytokines - CCL2, CCL3, CCL4, CCL5, CXCL10 and TNF - were elevated in patient sera, and were also robustly upregulated in microglia and OB lysates from GAS-infected mice (**Figure 2c–i; Extended Data Figure 2c**). GM-CSF was likewise significantly elevated in patient sera (**Table 1**). These findings indicate that the acute phase of PANDAS/PANS is associated with a systemic inflammatory signature overlapping with GAS-induced microglial responses in mice.

**Table 1.** Cytokines, chemokines and growth factors upregulated in sera from acute PANDAS/PANS patients.

| Serum protein | Healthy controls (pg/mL) | PANDAS/PANS (pg/mL) | p-value | p-adj |
| --- | --- | --- | --- | --- |
| CCL2 | 25.5 (8.475 - 105.5) | 157 (85.5 - 253.5) | 0.0005 | * |
| CCL3 | 4 (0.525 - 10.1) | 29 (14.5 - 47.5) | <0.0001 | ** |
| CCL4 | 81 (55.5 - 90.5) | 131 (93 - 181) | 0.0012 | * |
| CCL5 | 30 (15.25 - 41) | 296.5 (91.75 - 440.75) | <0.0001 | ** |
| CXCL8 | ND | 6.9 (2.1 - 27) | 0.0011 | * |
| CXCL10 | 7.2 (5.975 - 8.525) | 59 (32 - 78) | 0.0004 | ** |
| CXCL12 | 237 (176 - 313) | 664 (385.5 - 809) | 0.0007 | * |
| EGF | 16 (0.85 - 103) | 87 (40 - 152.5) | 0.014 | ns |
| GM-CSF | ND | 16 (4.075 - 24) | <0.0001 | ** |
| HGF | 54 (31.5 - 443) | 443 (358.5 - 589) | 0.0024 | ns |
| IL-1RA | ND | 536 (110.75 - 1115.75) | 0.0156 | ns |
| IL-6 | ND | 7.85 (2.05 - 26.5) | 0.0174 | ns |
| IL-7 | 1 (0.475 - 3.4) | 5.2 (3.55 - 7.65) | 0.0004 | ** |
| IL-12A/B | ND | 2.9 (1.95 - 3.85) | <0.0001 | ** |
| IL-15 | ND | 7.9 (3.525 - 24.5) | 0.0037 | ns |
| IL-22 | ND | 8.9 (4.175 - 30.5) | 0.0183 | ns |
| IL-23 | ND | 16 (5.5 - 44.5) | 0.0244 | ns |
| KITLG | 5.2 (3.45 - 7.4) | 14 (9 - 18) | 0.0009 | * |
| LIF | ND | 28 (18 - 58.5) | 0.0129 | ns |
| PDGFB | 87 (43.5 - 586.5) | 1800 (615.5 - 2620) | 0.0004 | ** |
| PGF | 94 (77 - 116) | 188 (96 - 290) | 0.0187 | ns |
| TNF | 5.2 (4.275 - 12.1) | 18 (9.975 - 22.5) | 0.0458 | ns |
| VEGFA | 88 (31.5 - 152) | 318 (138.5 - 483) | 0.0003 | ** |
| VEGFD | ND | 22 (1.65 - 32) | 0.0253 | ns |

### IFNγ derived from Th1 cells contributes to GAS-induced antigen presentation and interferon response in microglia and BECs in Th17-deficient mice

Elevated GM-CSF and other T cell - derived cytokines in patient sera (**Table 1**), together with prior evidence that Th17 cells drive GAS-induced BGE pathology [25] prompted us to examine how Th17 deficiency alters microglial and BEC responses. We performed scRNAseq on OBs from GAS-infected mice lacking the Th17 fate-specifying transcription factor RORγt [25, 56] (**Figure 4a**). These mice exhibit robust infiltration of IFNγ^⁺^Th1 cells, but markedly reduced Th17 cells after GAS infection [25]. BECs from GAS-infected *ROR*γ*t^-/-^* mice showed partial restoration of BBB-associated transcripts (e.g., *Mfsd2a*, *Tjp1* and *Itih5*), alongside increased expression of antigen-presentation and interferon response genes relative to wild-type (WT) mice (**Figure 4b; Supplementary Data Table 3**). Gene ontology (GO) pathway analysis confirmed enrichment of antigen presentation, interferon signaling and lymphocyte-related pathways, with concomitant downregulation of pathways related to vascular development and cell migration (**Figure 4c; Supplementary Data Table 5**).

**Figure 4.**
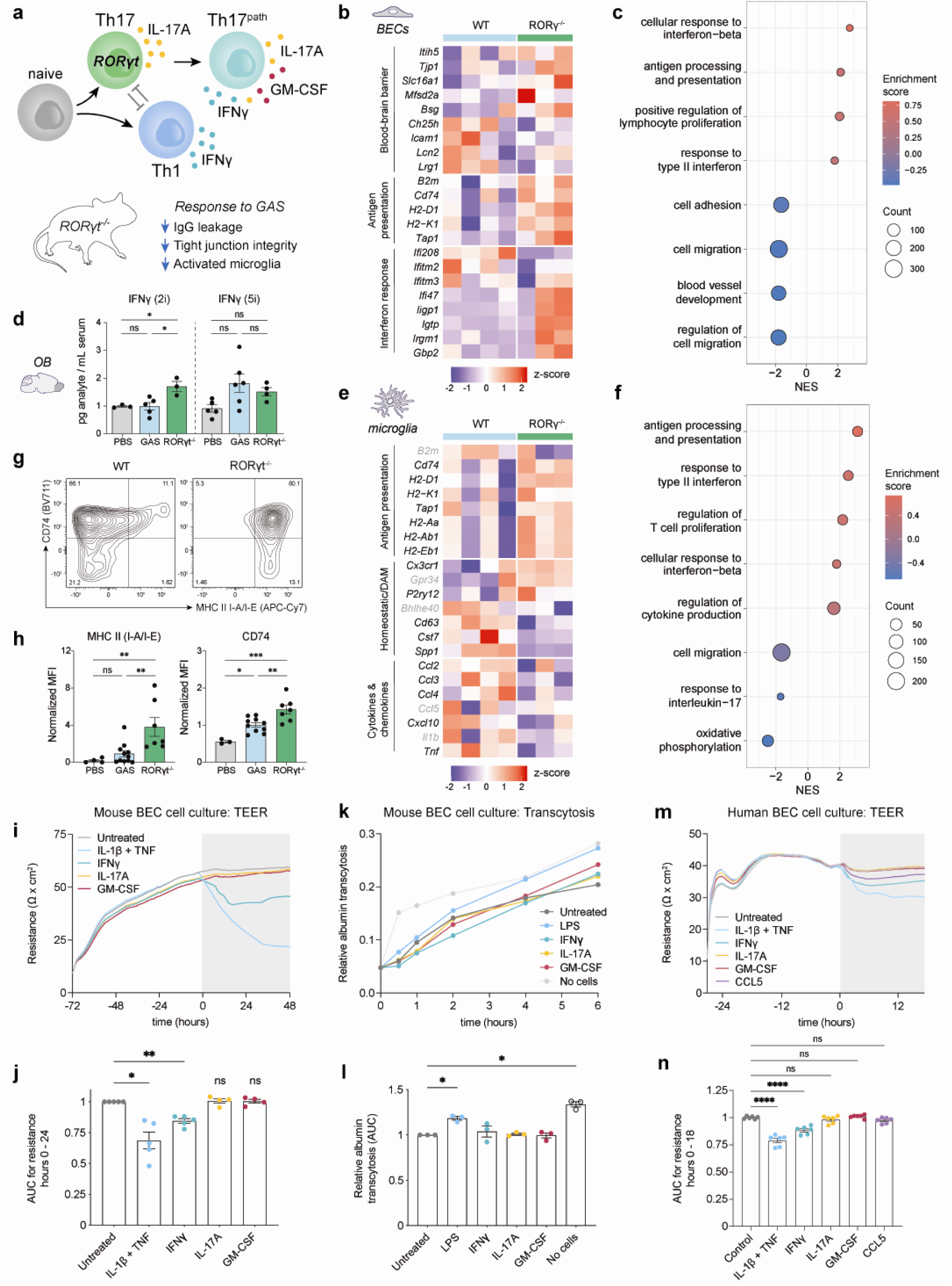
Th17-deficient mice show partial restoration of endothelial and microglial transcriptional responses after GAS infections. **a**, Schematic of CD4⁺ T cell differentiation, effector cytokine production, and associated brain pathology in *ROR*γ*t*^-/-^ mice following recurrent GAS infections. **b,** Heatmaps of differentially expressed genes (DEGs) related to BBB function, antigen presentation, and interferon response in olfactory bulb (OB) brain endothelial cells (BECs) from GAS-infected wild-type (WT) and *ROR*γ*t*^-/-^ mice. Values are shown as log(z-score); significantly altered genes are indicated in black (adjusted *p* < 0.05). **c,** Gene ontology (GO) pathway enrichment analysis of transcripts upregulated and downregulated in *ROR*γ*t*^-/-^ versus WT BECs after GAS infections. **d,** IFNγ concentrations in whole OB lysates from WT PBS (gray), WT GAS (blue), and *ROR*γ*t*^-/-^ GAS (green) mice after two or five infections. Statistical comparisons were performed using one-way ANOVA (ns, p > 0.05; *p* < 0.05). **e,** Heat maps of DEGs related to antigen presentation, homeostatic/DAM programs, and cytokine/chemokine signaling in OB microglia from GAS-infected WT and *ROR*γ*t*^-/-^ mice (log(z-score); adjusted *p* < 0.05 shown in black). **f,** GO pathway enrichment analysis of transcriptional changes in *ROR*γ*t*^-/-^ versus WT microglia following GAS infections. **g,** Representative flow cytometry plots of CD74 and MHC class II (I-A/I-E) expression in WT and *ROR*γ*t*^-/-^ microglia after GAS infections (n = 3,640 and 4,657 cells, respectively). **h,** Quantification of microglial surface expression of antigen presentation markers in WT PBS, WT GAS, and *ROR*γ*t*^-/-^ GAS mice. Statistical comparisons were performed using one-way ANOVA (ns, p > 0.05; *p* < 0.05; \**p* < 0.01). **i, j,** Transendothelial electrical resistance (TEER) measurements in primary mouse BEC monolayers following cytokine treatment, shown as representative ECIS traces (**i**) and area-under-the-curve (AUC) quantification (**j**). Gray shading indicates treatment period. Data are mean ± SEM (n = 5 replicates from 3 independent experiments). Mixed-effects analysis (*p* < 0.05; \**p* < 0.01). **k, l,** Relative transwell permeability of primary mouse BEC monolayers to albumin - AF594 following cytokine treatment. Untreated cells were used as reference, and LPS served as a positive control. Data are mean ± SEM (n = 3 independent experiments). Repeated-measures one-way ANOVA (*p* < 0.05). **m, n,** TEER measurements in primary human brain microvascular endothelial cells (HBMECs), shown as representative ECIS traces (**m**) and AUC quantification (**n**). Data are mean ± SEM (n = 6 replicates from 3 independent experiments). Mixed-effects analysis (\*\*\**p* < 0.0001).

Similarly, microglia from GAS-infected *ROR*γ*t^-/-^* mice displayed increased expression of antigen presentation genes, but reduced expression of multiple chemokines and cytokines (e.g. *Ccl2*, *Ccl3*, *Ccl4*, *Tnf*) (**Figure 4e; Supplementary Data Table 3**). GO analysis corroborated enrichment of antigen processing and immune response pathways in GAS-infected *ROR*γ*t^-/-^*microglia **(Figure 4f; Supplementary Data Table 5)**. Flow cytometry confirmed increased surface expression of CD74 and MHC class II (I-A/I-E) in microglia from GAS-infected *ROR*γ*t^-/-^*compared to WT mice (**Figure 4g, h**). Th17 cell-dependent suppression of antigen presentation genes extended beyond microglia and BECs, with increased MHC class I expression in astrocytes, OECs and neurons (**Extended Data Figure 5d**). A potential mechanism for this effect could be increased IFNγ in *ROR*γ*t^-/-^*mice, since IL-17A suppresses IFNγ expression and Th1 cell identity [57, 58]. IFNγ also increases MHC gene expression in myeloid cells and neurons [59, 60]. Consistently, IFNγ protein levels were significantly elevated in OBs from *ROR*γ*t^-/-^* mice after GAS infection (**Figure 4d**).

To understand how Th17-derived cytokines affect BBB function, we treated confluent primary mouse BECs and human brain microvascular endothelial cells (HBMECs) with either mouse or human IL-17A, GM-CSF or IFNγ, respectively. We measured the transendothelial electrical resistance (TEER) for 18-48 hours after cytokine treatment as a readout of paracellular barrier integrity. An IL-1β/TNF combination served as a positive control for this experiment since these cytokines are known to disrupt BBB integrity. *In vitro*, IFNγ, but not IL-17A or GM-CSF, reduced TEER in mouse and human BEC monolayers, indicating impaired paracellular barrier function, albeit to a lesser degree than IL-1β/TNF combination **(Figure 4 i, j, m, n)**. In addition, we also cultured mouse BECs to confluency in a transwell system and measured the transport of fluorescently labelled albumin from the top to the bottom chambers across the BEC monolayer as a proxy for transcellular barrier permeability. Although LPS (positive control) increased transcellular albumin transport across the mouse BEC monolayer, none of the Th17-derived cytokines showed any effect (**Figure 4k, l**). In summary, IFNγ induces antigen presentation and interferon responses in microglia and BECs, with modest direct effects on BBB integrity (**Figure 7a**).

### GM-CSF regulates a subset of microglial and BEC transcriptomic responses after multiple GAS infections

Since Th17 lymphocytes are required to induce brain pathology and a subset of transcriptome changes in microglia and BECs in our mouse model, we investigated the effects of Th17-derived cytokine (GM-CSF or IL-17A) on these transcriptome changes. To assess the role of GM-CSF, a cytokine downstream of RORγt implicated in autoimmune pathology [61], we first quantified GM-CSF–producing CD4^⁺^ T cells during GAS infection. The proportion of GM-CSF^⁺^CD4^⁺^ T cells and IFNγ^⁺^IL-17A^⁺^ (Th17^path^) cells increased in the OB with successive GAS infections (**Extended Data Figure 4a-c**). GM-CSF^+^ CD4^+^ T cells were reduced by 2-fold in *ROR*γ*t^-/-^* mice (**Extended Data Figure 4d**).

We next generated mice with CD4^⁺^ T cell–specific deletion of GM-CSF (*Csf2^ΔCD4^*) by crossing *CD4-Cre^ERT2^* [62] to *Csf2^fl/fl^* [63] mice and administering 4-OH-tamoxifen daily between P16 and P20 prior to onset of GAS infections at P28 (**Extended Data Figure 4e**). This protocol resulted in a significant decrease in GM-CSF^+^ CD4^+^ T cells (**Extended Data Figure 4f**). Csf2^ΔCD4^ mice showed normal survival and CD4^⁺^ T cell infiltration after GAS infection **(Extended Data Figure 4g, h)**. scRNAseq revealed that microglia from *Csf2^ΔCD4^* mice exhibited reduced expression of cytokine and chemokine transcripts, with minimal changes in antigen presentation genes compared to *Csf2^fl/fl^*microglia after GAS infections (**Extended Data Fig. 4i; Supplementary Data Table 3**). GO analysis confirmed downregulation of cytokine and interferon response pathways (**Extended Data Figure 4j; Supplementary Data Table 5)**. Despite these transcriptomic changes, CD68^⁺^Iba1^⁺^ microglia numbers were increased (**Extended Data Fig. 4k, l**), and CD74 protein expression was unchanged in *Csf2^ΔCD4^* mice (**Extended Data Figure 4m, n**), although this phenotype was not accompanied by changes in cell cycle related transcripts (**Supplementary Data Table 3**).

In BECs, GM-CSF deletion resulted in modest upregulation of select BBB-related genes (e.g. *Itih5*, *Bsg*) and reduced expression of antigen presentation, response to bacterium and interferon-response pathways corroborated by GO analysis **(Extended Data Figure 4o, p; Supplementary Data Table 3, 5).** Consistent with scRNAseq data, there was no improvement in BBB permeability to serum IgG in the *Csf2^ΔCD4^* OB after GAS infections **(Extended Data Figure 4q-s)**. Together, these findings indicate that CD4^+^ T cell-derived GM-CSF preferentially modulates microglial inflammatory programs, with limited impact on BBB integrity.

### Global IL-17A blockade reverses GAS-induced transcriptome changes in BECs and microglia, but exacerbates BBB dysfunction during active infection

To assess the role of IL-17A, a major cytokine produced by Th17 cells, mice were treated with an IL-17A–neutralizing antibody or isotype control throughout GAS infection (**Figure 5a**). IL-17A blockade did not alter CD4^+^ T cell infiltration, or subtype distribution in the OB (**Extended Data Figure 5a-c**). In BECs, IL-17A neutralization increased expression of select BBB-associated genes, particularly regulators of transcytosis, as well as antigen presentation, and shifted GO pathways toward metabolic and maturation-associated BEC programs such as cellular respiration and oxidative phosphorylation **(Figure 5b, c, Supplementary Data Table 5)**. Conversely, inflammatory and interferon-related pathways were downregulated after IL-17A blockade **(Figure 5b, c, Supplementary Data Table 5)**. Despite these transcriptomic changes, IL-17A blockade worsened BBB permeability to serum IgG **(Figure 5d-f)**, accompanied by further reductions in *Itm2a* mRNA and tight junction proteins, Claudin-5 and ZO-1 **(Figure 5k-m)**. Although, a subset of interferon response genes (e.g. *Ifit1,2,3*, *Ifitm2,3*) were rescued at the scRNAseq level (**Supplementary Data Table 3)**, there was no change in IFITM3 protein level in OB CD31^+^ blood vessels between two treatments after GAS infections (**Figure 5g-i**). IL-17A is critical to fight GAS infections in the periphery [64], therefore its blockade could increase peripheral inflammatory responses. Serum cytokine levels including IFNγ, CCL4, CCL5, CXCL2, CXCL10 and GM-CSF, were elevated in IL-17A - treated mice **(Figure 5n),** and survival rate after GAS infections was reduced **(Figure 5j)**, consistent with impaired control of the peripheral infection.

**Figure 5.**
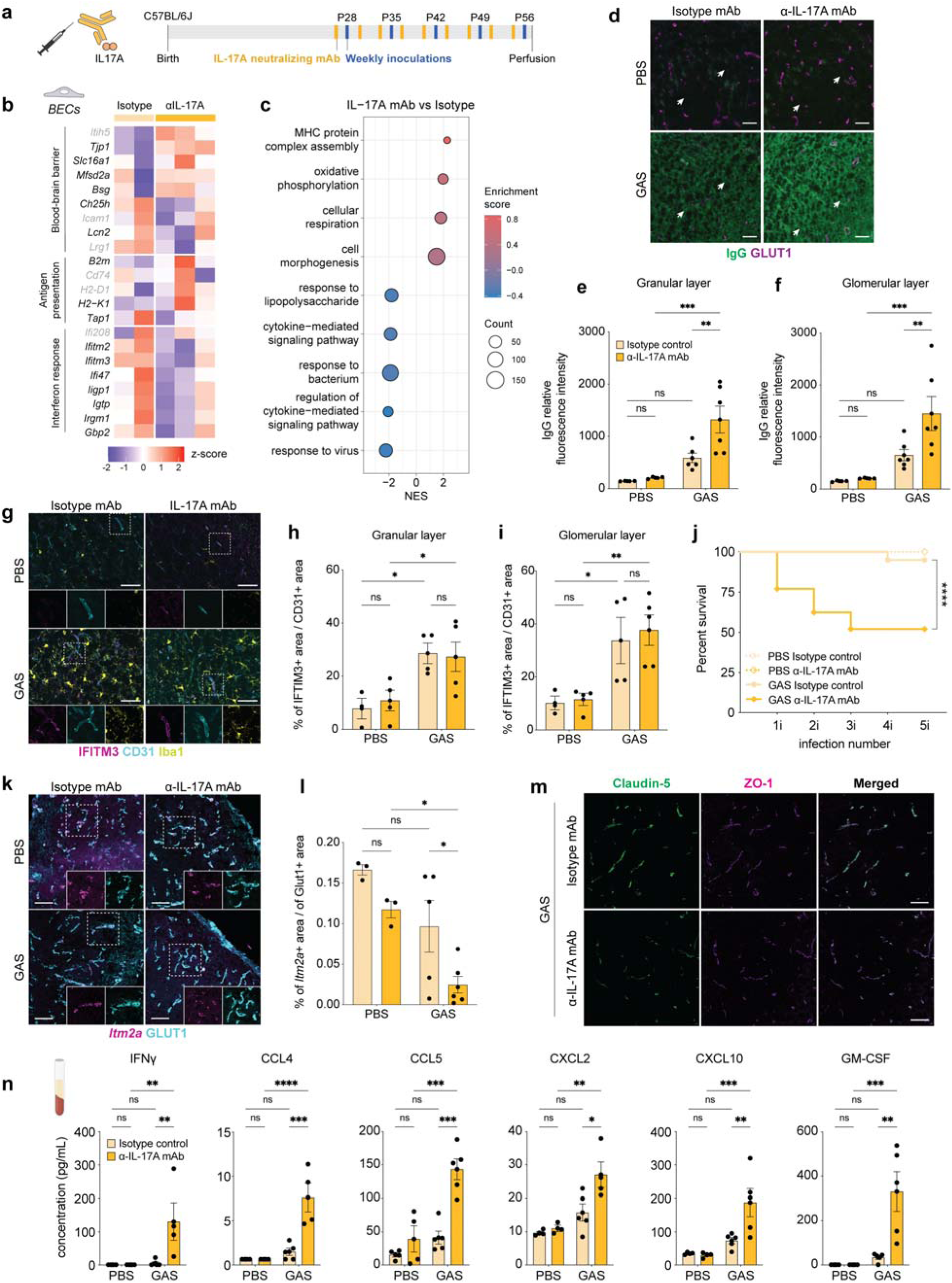
IL-17A inhibition partially restores endothelial transcriptional programs after GAS infections but worsens BBB dysfunction *in vivo*. **a,** Timeline of recurrent intranasal GAS infections and administration of an α–IL-17A–neutralizing monoclonal antibody (mAb) or isotype control. **b,** Heat maps of differentially expressed genes (DEGs) related to BBB function, LPS response, interferon signaling, and antigen presentation in olfactory bulb (OB) brain endothelial cells (BECs) from GAS-infected mice treated with isotype control or α–IL-17A mAb. Values are shown as log(z-score); significantly altered genes are indicated in black (adjusted *p* < 0.05). **c,** Gene ontology (GO) pathway enrichment analysis of transcripts upregulated and downregulated in α–IL-17A mAb–treated versus isotype-treated BECs following GAS infections. **d,** Representative immunofluorescence (IF) images of endogenous IgG leakage (green) in the granular layer of the OB. Blood vessels are labeled with Glut1 (cyan). **e, f,** Quantification of IgG extravasation (relative fluorescence intensity) in the glomerular (**e**) and granular (**f**) OB layers from PBS- or GAS-infected mice treated with α–IL-17A mAb or isotype control. Comparisons were performed using two-way ANOVA with Šídák’s multiple comparisons test (*p* < 0.05; \*\**p* < 0.001; n = 3–7 mice per group). **g–i,** Representative IF images for IFITM3 (pink), Iba1 (yellow), and CD31 (blue) in the OB of GAS-infected mice treated with isotype or α–IL-17A mAb (**g,h**), and quantification of Ifitm3-positive area within CD31⁺ vasculature (**i**). **j,k,** Representative fluorescence in situ hybridization (FISH) images of *Itm2a* mRNA (pink) combined with Glut1 (blue) in the glomerular layer (**j**) and quantification of vascular Itm2a signal (**k**). **l,** Representative IF images of the tight junction proteins Claudin-5 (green) and ZO-1 (red) in the OB vasculature of GAS-infected mice treated with isotype or α–IL-17A mAb. **m,** Serum cytokine concentrations measured by multiplex immunoassay in PBS- or GAS-infected mice treated with isotype or α–IL-17A mAb. Data are mean ± SEM. Comparisons were performed using one-way ANOVA with Tukey’s multiple comparisons test (ns, p > 0.05; *p* < 0.05; \**p* < 0.01; \*\**p* < 0.001; \*\*\**p* < 0.0001; n = 3 - 6 mice per group). **n,** Survival curves of GAS-infected mice treated with isotype control or α–IL-17A mAb (n = 13–48 mice per group). Statistical significance was assessed using the Mantel–Cox log-rank test.

Microglial scRNAseq and GO analysis from the IL-17A mAb-treated condition revealed partial restoration of homeostatic gene expression and reduced DAM-associated transcripts and cytokine production, but increased antigen presentation and interferon-response pathways paralleling the transcriptome phenotype of *ROR*γ*t^-/-^* microglia (**Extended Data Figure 5d, e; Supplementary Data Table 3, 5**). Genes related to antigen presentation by MHC class I were also upregulated in other OB cell types, including OECs and astrocytes after IL-17A blockade (**Extended Data Figure 5f**). In contrast to the *ROR*γ*t^-/-^*microglial phenotypes observed by IF [25], the number of activated CD68^+^ Iba1^+^ myeloid cells remained unchanged after IL-17A mAb treatment, and there were was no difference in microglial expression of CD74 and IFITM3 proteins between the two treatments after GAS infections (**Extended Data Figure 5g-l**), likely reflecting heightened peripheral inflammation. These findings suggest that systemic IL-17A blockade during active infection exacerbates neurovascular pathology through indirect peripheral immune effects.

### Microglia-specific ablation of IL-17 receptor A rescues BBB dysfunction without altering microglia activation

To determine whether IL-17A signaling within the brain contributes directly to pathology, we examined IL-17 receptor expression and found high IL17RA expression in microglia, macrophages and neutrophils **(Extended Data Figure 3d)**, suggesting a critical contribution of these cells to pathology. To address this hypothesis, we generated mice lacking IL17RA in microglia/macrophages (*Il17ra ^ΔCX3CR1^*) by crossing *CX3CR1^CreERT2^* [65] with *Il17ra^fl/fl^*[66] mice and administering 4-OH tamoxifen between P16 and P20 prior before GAS infections at P28 **(Figure 6a)**. IL17RA deletion in microglia did not significantly alter CD4^+^ T cell infiltration into the OB **(Extended Data Figure 6a, b)**, but significantly reduced BBB permeability to serum IgG in the granular layer **(Figure 6b-d)**. Consistent with a partial rescue in BBB function, Mfsd2a protein levels showed a non-significant increase in the OB vasculature **(Figure 6g, h)**, whereas *Itm2a* mRNA expression and tight junction markers (Claudin-5 and ZO-1) were comparable between genotypes after GAS infections (**Figure 6e, f, j**). The vascular expression of IFITM3 protein was also increased, although not significantly, in the OB glomerular layer in *Il17ra ^ΔCX3CR1^* mice after GAS infections **(Figure 6i; Extended Data Figure 6g)**. Importantly, *Il17ra ^ΔCX3CR1^* mice did not exhibit reduced survival after GAS infection **(Figure 6k),** in contrast to systemic IL-17A blockade. Finally, *Il17ra ^ΔCX3CR1^* mice showed no difference in either microglial activation (CD68^+^ Iba1^+^), expression levels for the antigen presentation marker CD74, or IFITM3 protein compared to *Il17ra^fl/fl^* GAS-infected mice **(Extended Data Figure 6c-h)**. Therefore, IL-17A/IL17RA signaling in microglia contributes to BBB dysfunction after GAS infection, while other aspects of microglial activation are likely maintained by other T cell - derived cytokines.

**Figure 6.**
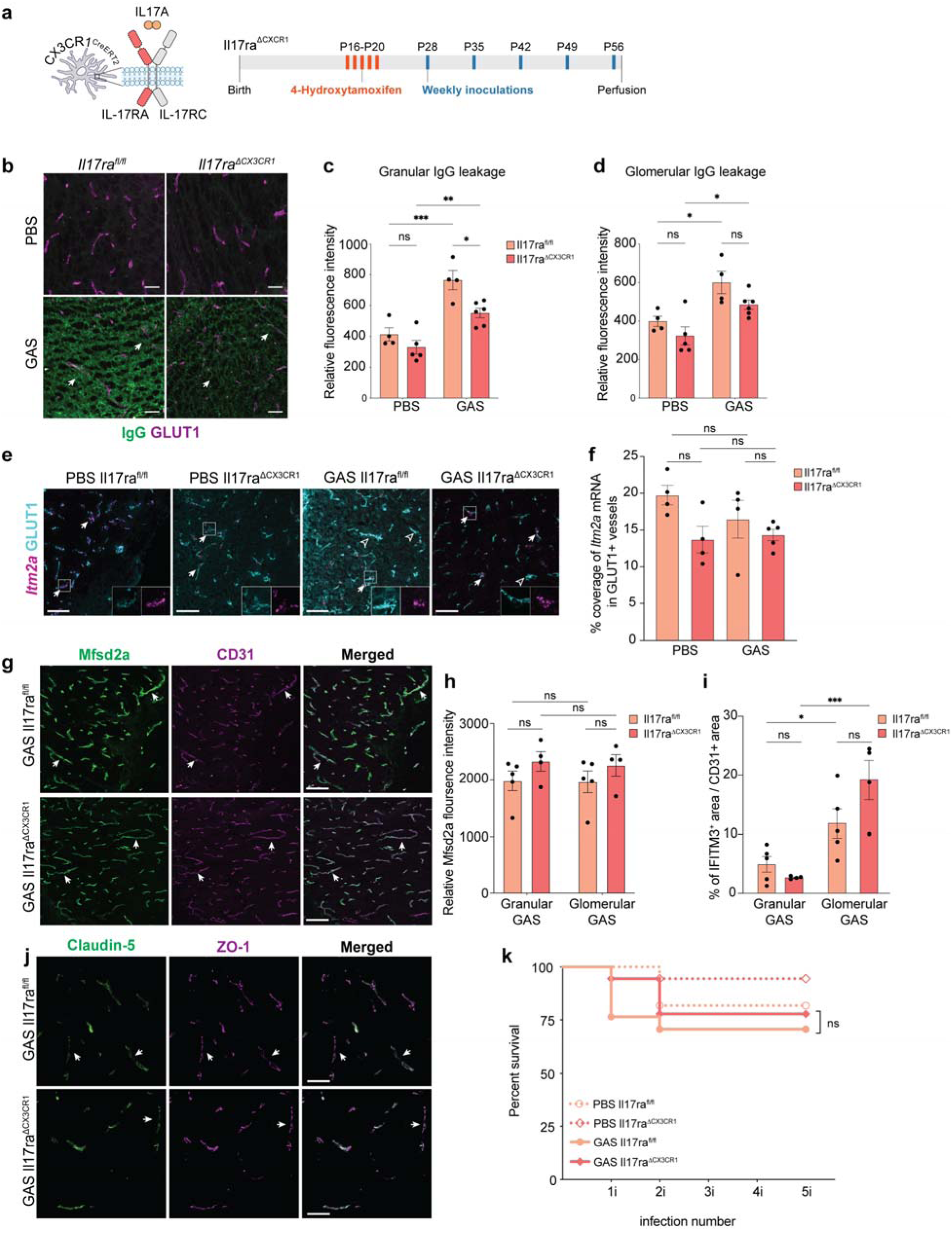
Myeloid-specific deletion of IL-17RA partially mitigates BBB disruption following recurrent GAS infections. **a,** Timeline of 4-hydroxytamoxifen (4-OHT) administration and recurrent GAS infections in *Il17ra^fl/fl^* (control) and *Il17ra*^Δ*CX3CR1*^ mice. **b–d,** Representative immunofluorescence (IF) images of endogenous serum IgG leakage (green) and Glut1-labeled vasculature (magenta) in the olfactory bulb (OB), and quantification of IgG extravasation in the glomerular and granular OB layers from PBS- or GAS-infected *Il17ra^fl/fl^*(salmon) and *Il17ra*^Δ*CX3CR1*^ (red) mice. **e, f,** Representative fluorescence in situ hybridization (FISH) images of *Itm2a* mRNA combined with Glut1 (BEC marker) in the glomerular OB layer, and quantification of vascular *Itm2a* signal in PBS- and GAS-infected mice of both genotypes. **g, h,** Representative IF images of Mfsd2a (green) and CD31 (purple) in the OB vasculature of GAS-infected *Il17ra^fl/fl^* and *Il17ra*^Δ*CX3CR1*^ mice, with quantification of Mfsd2a mean fluorescence intensity (MFI). **i,** Quantification of IFITM3-positive area within CD31⁺ endothelial regions in the OB from GAS-infected mice of both genotypes. Graph bars represent mean ± SEM. Comparisons were performed using two-way ANOVA with Šídák’s multiple comparisons test (ns, p > 0.05; **p** < 0.01; \***p** < 0.001; \*\***p** < 0.0001; n = 4–8 mice per group). **j,** Representative IF images of the tight junction proteins Claudin-5 (green) and ZO-1 (red) in the glomerular layer of the OB from GAS-infected mice of each genotype. **k,** Survival curves of PBS- or GAS-infected *Il17ra^fl/fl^* and *Il17ra*^Δ*CX3CR1*^ mice (n = 11–18 mice per group). No significant differences were observed between GAS-infected genotypes (Mantel–Cox log-rank test).

## Discussion

GAS infections are a major cause of morbidity in children and adults worldwide [67], not only due to primary infections such as pharyngitis, but also through secondary neuropsychiatric sequelae [4]. While aberrant cellular and humoral immune responses underlie these CNS complications (reviewed in [3–5]), the molecular changes in brain cells, and the contributions of T cell-derived cytokines are not understood. Using genetic and antibody blockade approaches in mice together with scRNAseq, spatial transcriptomics, and validation experiments, we mapped the transcriptional changes in CNS cells - particularly BECs and microglia - following multiple GAS infections. We find that microglia in close proximity to infiltrating CD4^⁺^ T cells exhibit the most robust “Streptococcus-responsive” gene expression. We further dissect shared and unique effects of two Th17 effector cytokines, IL-17A and GM-CSF, on microglial and BEC transcriptomes. Finally, serum from PANDAS/PANS patients displays elevated levels of multiple inflammatory cytokines, highlighting a potential role for Th17 lymphocytes in human disease progression.

### Transcriptional remodeling of BECs and microglia

scRNAseq of over 100,000 OB cells revealed that BECs and microglia are among the CNS cell types most transcriptionally altered after GAS infections. Both upregulate inflammatory programs, including antigen presentation, interferon response, and cytokine production, while downregulating homeostatic signatures: BBB-associated genes in BECs and homeostatic genes in microglia. BEC transcriptional responses mirror patterns seen in other inflammatory conditions, such as systemic LPS administration [39, 68, 69], and EAE [70], with upregulation of antigen presentation, interferon response, and downregulation of BBB-associated transcripts [40]. These molecular changes are consistent with our prior findings that BBB structure and function are impaired after GAS infections [23, 25]. Despite increased BBB permeability to large proteins (e.g. IgG), genes promoting caveolar transport (e.g., *Cav1*, *Cav2*, *Cavin1–2*) were downregulated, suggesting that decreased Mfsd2a expression contributes to elevated transcellular transport [43]; however, junctional breakdown or bulk transcytosis may also play roles. *Lcn2*, strongly induced in BECs, may mediate IL-17A-dependent innate immune responses, consistent with its role in other diseases [41, 42].

Microglia similarly exhibit upregulation of DAM, interferon response, antigen presentation, and cytokine/chemokine genes. Heterogeneity exists among the four Streptococcus-responsive microglia clusters, with srMG4 displaying high interferon-response genes and downregulation of some classic DAM genes (e.g., *Mertk*, *Trem2*), indicating an activation state distinct from that seen in neurodegeneration models [71]. T cells that infiltrate the brain after GAS infections secrete IFNγ (type II interferon), and we could not detect IFNα or -β (type I interferons) either transcriptionally or by multiplex cytokine immunoassay in the OB (data not shown). Microglial expression of interferon-response genes has been seen in inflammation [72, 73], autoimmunity [74–76], injury [77, 78], neurodegeneration [79–81] and aging [82] and is implicated in cell morphology changes, inflammation and phagocytosis, among other microglial functions [82–84]. IFNγ produced by infiltrating T cells appears to drive antigen presentation gene expression in microglia, BECs, and other CNS cells after GAS infections (**Figure 7a**), consistent with known effects of type II interferons on MHC regulation. Upregulated microglial chemokines (e.g., CCL2, CCL3, CCL4, CCL5, CXCL10) may facilitate recruitment of peripheral immune cells, and combined with T cell effector cytokines, could impair BBB integrity *in vivo*. For example, CCL2-CCR2 signaling is required for Th17^path^ cell recruitment to the CNS in EAE and *S. pneumoniae* infections [85]. Several microglia-derived cytokines (e.g. TNF, IL-1β) and chemokines (e.g. CCL2, CCL5) are known to break down the BEC barrier *in vitro*, and they may contribute, together with T cell effector cytokines, to impair BBB function *in vivo*. MERFISH analysis confirms that microglial transcriptional shifts are greatest in OB regions with dense T cell infiltration, supporting a key role for T cell - microglia crosstalk in driving CNS pathology as predicted by Cell Chat.

**Figure 7.**
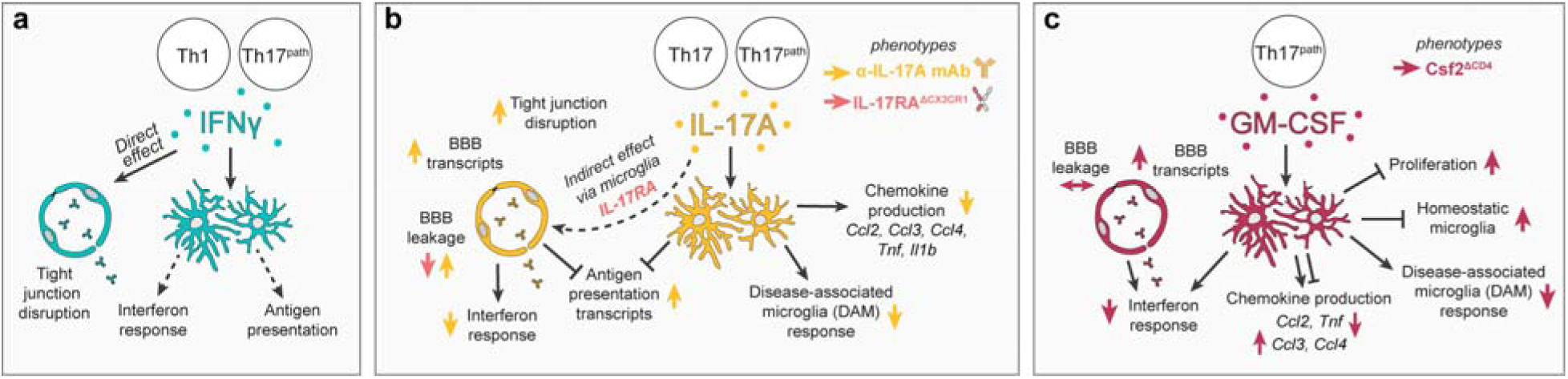
Model summarizing the putative effects of IFNγ, IL-17A, and GM-CSF on brain endothelial and microglial responses to recurrent GAS infections. **a–c,** Proposed model illustrating how effector cytokines produced by infiltrating Th1 and Th17 lymphocytes following repeated GAS infections may differentially shape transcriptional responses in brain endothelial cells (BECs) and microglia. **a,** IFNγ is predicted to contribute to the induction of interferon-response programs and antigen-presentation pathways in both microglia and BECs. **b,c,** IL-17A and GM-CSF appear to exert partially overlapping effects on microglial activation, including regulation of proliferative responses and induction of disease-associated microglial (DAM) genes, cytokines, and chemokines, while also displaying cytokine-specific contributions. Notably, IL-17A signaling may more strongly influence endothelial transcriptional alterations following GAS infections. In addition, antigen-presentation signatures remain elevated in conditions of Th17 deficiency or systemic IL-17A neutralization, suggesting that IL-17A may normally modulate these pathways in both microglia and BECs. Together, these models highlight shared and distinct roles for Th17-associated cytokines in shaping neuroimmune and vascular responses after recurrent GAS exposure.

### Th17 cytokines drive distinct transcriptomic responses

The transcriptomic analysis of microglia and BECs from *ROR*γ*t*^-/-^ mice supports our prior findings that Th17 cells are critical for brain pathology in our mouse disease model [25]. Loss of Th17 cells partially rescues microglial chemokine/cytokine expression and BEC BBB transcripts while paradoxically increasing antigen presentation and interferon-response genes, likely due to elevated IFNγ levels and absence of IL-17A/GM-CSF inhibition. IFNγ is known to increase MHC gene expression in myeloid cells and neurons [59, 60], and its production by T cells can be inhibited by IL-17A [57, 58]. Circulating IFNγ levels may be responsible for upregulation of antigen presentation and the interferon response in microglia and BECs (**Figure 7a**).

IL-17A and GM-CSF have overlapping and distinct effects on microglia. Both contribute to microglial DAM gene expression and cytokine/chemokine regulation, but specific chemokine subsets are preferentially regulated by one cytokine (**Figure 7b, c**). IL-17A more effectively suppresses antigen presentation in microglia and BECs, as evidenced by increased expression in both *ROR*γ*t^-/-^* and IL-17A mAb-treated mice. In contrast while interferon response genes are strongly downregulated in microglia from *Csf2^ΔCD4^*, *ROR ^--/-^* and IL-17A mAb-treated mice, expression of specific chemokine transcripts seems to be preferentially regulated by either IL-17A or GM-CSF (**Figure 7b, c**). In contrast, the effects of IL-17A and GM-CSF are quite divergent in BECs. GM-CSF deficiency in T cells minimally affects BBB transcriptome or permeability, but reduces inflammatory signaling in microglia, highlighting a predominantly microglia-targeted effect. IL-17A blockade rescues transcriptomic shifts in BECs but worsens BBB permeability, likely due to unresolved peripheral inflammation. IL-17A does not act directly on BECs, because it cannot disrupt tight junctions, nor promote albumin transcytosis in primary cultured mouse or human BECs (**Figure 4i-n**) consistent with robust IL17RA expression in microglia and macrophages, but not BECs, after GAS infections. Genetic ablation of IL17RA in microglia/macrophages partially rescues BBB leakage, indicating microglia mediate IL-17A effects on the vasculature (**Figure 7d**). Future studies will identify downstream effectors of IL17A/IL17RA signaling in microglia and how they affect BBB function after GAS infections.

### Clinical relevance

Currently, it is contested whether PANDAS/PANS has an inflammatory origin, although neuroinflammation has been seen in the basal ganglia of PANDAS patients [9] using a PET ligand that binds both activated microglia and astrocytes [86]. PANDAS/PANS patients exhibit elevated serum cytokines, chemokines, and growth factors during acute disease, mirroring cytokine upregulation in GAS-infected mouse OBs. Some elevations, including IL-6 and CXCL10, are shared with acute GAS pharyngitis cases [87], whereas patients with invasive GAS infections have elevated levels of IL-1β, IL-6, IL-8, IL-10 and IL-18 [88]. Other elevated cytokines (CCL2, CCL4, CCL5, IL-7 and TNF) appear specific to neuropsychiatric sequelae, supporting a distinct inflammatory profile. Distinguishing PANDAS/PANS from Tourette’s syndrome (TS) or OCD remains challenging as overlapping cytokine elevations are reported. Elevated levels of IL-17A and TNF have been found in sera from PANDAS/PANS patients [53, 54] and high TNF and IL-12 serum levels correlate with symptom exacerbation in children with tic disorders and OCD [89, 90]. In addition, immune phenotypes linked to TS include elevated levels of IL-6, IL-12, IL-17A and TNF [89, 91]. Our data provide new candidate biomarkers for acute SC/PANDAS, which may aid diagnosis and clarify the role of neuroinflammation in these disorders.

## Acknowledgements

We thank Michael Kissner and Rosemary Gordon-Schneider from the Columbia Stem Cell Initiative Flow Cytometry Core for technical support; Erin Bush and Izabela Krupska from the Single Cell Analysis Core, JP Sulzberger Columbia Genome Center (CUIMC) for library construction and sequencing; Dr. Wassim Elyaman for sharing *TMEM119-tdTomato* and *CX3CR1-GFP* mouse lines; S.V. Pollak from the Biomarkers Core Lab, Irving Institute for Clinical and Translational Research (CUIMC) for multiplex cytokine analysis; Drs. Ai Yamamoto (CUIMC) and Arnold Han (CUIMC) for equipment use; Dr. Ilir Agalliu (Albert Einstein College of Medicine) for advice on statistical analyses; Dr. Chenghua Gu (Harvard Medical School) for one of the Mfsd2a antibodies used in the study; Dr. Alfred Simkin for the find_nearest_neighbors2 python script; and Drs Jennifer Bain, MD (CUIMC), Wendy Silver, MD (CUIMC), Jay Selman, MD (CUIMC), Rebecca Hommer, MD (NIMH) and Hannah Z. Street (CUIMC) for help with patient recruitment and sample collection.

The work in the Agalliu laboratory was supported in part by NIMH (R01MH112849 and R56MH109987); NINDS 2T32NS064928 (CRW); NHLBI (R61/R33 HL159949) and NIA (RF1AG078352), NEI (R01EY033994), International OCD Foundation (DA, UA), PANDAS Network (UA), PANDAS Physician Network (UA) and the Global Lyme Alliance (DA, UA) and National Center for Advancing Translational Sciences, National Institutes of Health, through Grant Number UL1TR001875 (UA). The work in the Schafer laboratory was supported by NIMH (R01MH113743, NINDS (R01NS117533), NIA (RF1AG068281 and R01AG068281), Massachusetts Life Sciences Center, BrightFocus Foundation A2022006F, Alzheimer’s Association AARF-22-923219, and the Dr. Miriam and Sheldon G. Adelson Medical Research Foundation.

We are grateful to the following major donors for their generous support of this research: Tom and Patti Walz, Newport Equities LLC, PANDAS Network, Northwest PANDAS/PANS Network, Steve and Wendy Swyter, the Look Foundation, The Alex Manfull Foundation. We also acknowledge the many families whose children were affected by this disease for their contributions to this study.

## Author information

Conceptualization: CRW, TC, DA; Animal experiments: CRW, UA, DA; RNA sequencing experiments: CRW, VM; MERFISH experiments and analysis: VDL, TEF, CRW, DPS; Bioinformatic analyses: CRW, TEF, VM; Flow cytometry experiments: CRW, LB, SJH; Immunofluorescence experiments: CRW, UA, LB, SJH, DA; *In situ* hybridization experiments: DA, BA, CRW; Data analyses: CRW, UA, DJ, DA; Cell culture experiments: CRW, PAF, TC, DA, NA, PA; Patient history & sample collection: SLD, SS, WV; Processing and analysis of patient samples: TC, NA, PA; Statistical analyses: CRW; Resources: DA, BC, DPS, RC, WE; Funding acquisition: DA, TC, UA; Supervision: DA, TC, DPS; Writing: CRW, DA; Revising: CRW, UA, TC, DA.

## Ethics declarations

None

## Online Methods

### Mice

Experiments involving mice were approved by Columbia University Irving Medical center (CUIMC) Institutional Animal Care and Use Committees. Mice were bred in the CUIMC vivarium, under 12-hour light/12-hour dark, pathogen-free conditions. Female mice were used for all experiments, except the time course analysis of Th17 cell subtypes by flow cytometry and *Csf2* recombination confirmation flow cytometry, which used even numbers of males and females. The *ROR*γ*t^eGFP^*mice[25, 56] (B6.129P2(Cg)-Rorctm2Litt/J, strain 007572) were obtained from the Jackson Laboratory. The *Csf2^fl/fl^* mouse strain [63] was provided by Bogoljub Ciric (Thomas Jefferson University, Philadelphia, PA). *CD4-CreER^T2^* transgenic mice (B6(129X1)-Tg(Cd4-cre/ERT2)11Gnri/J, strain 022356) [62] were obtained from the Jackson Laboratory and crossed to *Csf2^fl/fl^*mice for two generations. *CD4-CreER^T2+/-^ Csf2^fl/fl^*males were mated to *Csf2^fl/fl^* females to generate *CD4-CreER^T2+/-^ Csf2^fl/fl^* (*Csf2^ΔCD4^*) experimental mice and *Csf2^fl/fl^*littermate controls. *CX3CR1-CreER^T2^* transgenic mice (B6.129P2(Cg)-*Cx3cr1^tm2.1(cre/ERT2)Litt^*/WganJ, strain 021160) [65] were obtained from the Jackson Laboratory and crossed to *Il17ra^fl/fl^* mice (B6.Cg-*Il17ra^tm2.1Koll^*/J, strain 03100) [66] for two generations. *CX3CR1-CreER^T2+/-^ Il17ra^fl/fl^* males were mated to *Il17ra^fl/fl^* females to generate *CX3CR1-CreER^T2+/-^Il17ra^fl/fl^* (*Il17ra^ΔCX3CR1^*) experimental mice and *Il17ra^fl/fl^* littermate controls. P16 pups were intraperitoneally injected daily with 100 µg of (Z)-4-Hydroxytamoxifen (Millipore Sigma, H7904), dissolved in 50 µL of corn oil (Millipore Sigma, C8267) for 5 days (P16-P20). *TMEM119-tdTomato* [47] and *CX3CR1-GFP* [46] reporter mouse lines were provided by Dr. Wassim Elyaman (CUIMC).

### Human serum studies

The experiments with human sera were approved by CUIMC Institutional Review Board. The NIMH sera used in this study were analyzed in a previous publication [92], and were obtained from the NIMH. Informed consent / assent was obtained from all subjects both at NIMH and CUIMC.

### GAS intranasal infections

Mice received weekly intranasal inoculations with either a suspension of *Streptococcus pyogenes* [Group A *Streptococcus* (GAS)], or phosphate-buffered saline (PBS) control, starting at P21-P28. Recombinant GAS strain expressing a 2W epitope-tagged M protein was used for all experiments as described [22, 23, 25]. GAS was streaked out on new blood agar plates each week. Culture media consisted of an autoclaved solution of 3% Todd-Hewitt Broth (Bacto, 90003-430) and 2% Neopeptone (Bacto, 90000-268). Several GAS colonies were used to inoculate 10 mL of culture medium and incubated overnight at 37°C in 5% CO_2_. The following day, the culture was diluted to an OD_600_ of 0.2 and grown to OD_600_ 0.6, centrifuged and washed in 1 mL PBS (without Ca^2+^ and Mg^2+^) and resuspended in 110 µL of PBS. The GAS suspension was kept briefly on ice prior to intranasal infections. All intranasal infections were performed in an ABSL2 vivarium facility. Mice were immobilized with light anesthesia and a P20 pipette was used to drip GAS suspension into nostrils. To reduce lethality due to sepsis, a smaller GAS dose was used during the first two weeks (the first GAS inoculation is 8 x 10^7^ CFU per nostril, the second is 12 x 10^7^ CFU per nostril, and the third, fourth and fifth are 2 x 10^8^ CFU per nostril). During the first two weeks of GAS infections, mice were provided with nutritional supplements (DietGel and ClearH_2_O, 72-27-5022).

### Neutralizing antibody treatment

Starting 24 hours prior to the first GAS infection, mice were injected intraperitoneally twice weekly with 500 µg of either InVivoMAb anti-mouse IL-17A monoclonal antibody, clone 17F3 (Bio X Cell catalog, BE0173), or mouse IgG1 isotype control monoclonal antibody, clone MOPC-21 (Bio X Cell, catalog BE0083), in 100 µL of dilution buffer (Bio X Cell catalog IP0070 and IP0065, respectively).

### Single-cell RNA sequencing

Mice were anesthetized with isoflurane and perfused intracardially with PBS for 3 minutes. Nasal associated lymphoid tissue (NALT), olfactory epithelium (OE), or olfactory bulb (OB) were dissected and placed in Hanks’ Balanced Salt Solution (HBSS) without Ca^2+^ and Mg^2+^ and cut up with a sterile scalpel blade. Two or three animals were pooled per sample. Tissue was then placed in C Tubes (Miltenyi Biotec, 130-093-237), along with dissociation reagents from the MACS Neural Tissue Dissociation Kit (P) (Miltenyi Biotec, 130-092-628). Samples were loaded onto a gentleMACS Octo Dissociator with Heaters (Miltenyi Biotec, 130-096-427) and the 37C_NTDK_1 program was run. Following dissociation, samples were filtered through a 70 µm cell strainer, washed in HBSS and resuspended in MACS buffer with myelin removal beads (Miltenyi Biotec, 130-096-733), then purified with an LS column (Miltenyi Biotec, 130-042-401), according to manufacturer instructions. Eluent was washed twice, incubated with DRAQ5 (Bio-Legend, 424101, 1:1,000) and CD16/CD32 Fc block (BD Biosciences, 553141, 1:200) at room temperature for 15 minutes. Cells were washed and incubated with antibodies against CD31 (FITC, BD Biosciences, 561813, 1:200) and CD11b (BV421, BioLegend, 101235, 1:100) for 30 minutes on ice. Cells were washed and resuspended in FACS buffer with propidium iodide (1:10,000). Live, nucleated cells (DRAQ5^+^ PI^lo^) were sorted on a FACSAria II (BD), equipped with 355 nm, 405 nm, 488 nm, 561 nm and 640 nm lasers and a 130 µm nozzle. In a subset of experiments, CD31^+^ and CD11b^+^ populations were collected to enrich for cell types of interest. Sequencing was performed by the Columbia Single Cell Core using 10X Genomics Chromium Single Cell 3’ technology, with reads aligned to the mm10-2020-A transcriptome.

### Immunofluorescence

Mice were anesthetized with isoflurane and perfused intracardially with PBS for 4 minutes, followed by 4% paraformaldehyde (PFA) for 6 minutes. Brains were extracted and post-fixed in 4% PFA for 4-6 hours, then washed three times in PBS, incubated overnight in 30% sucrose, embedded in Tissue-Plus O.C.T. compound (Fisher, 4585) and stored at -80°C. Coronal sections (12 µm) were cut on a Leica CM3050 S Cryostat and stored at -80°C. For immunofluorescence staining, slides were washed in PBS for 10 minutes, incubated for 1 hour at room temperature in blocking buffer (10% BSA in 1X PBS with 0.1% Triton-X-100), and with primary antibodies diluted in PBST (0.1% Triton-X-100 in 1X PBS) with 1% BSA overnight at 4°C. The table below provides a complete list of immunofluorescence antibodies used for this study.

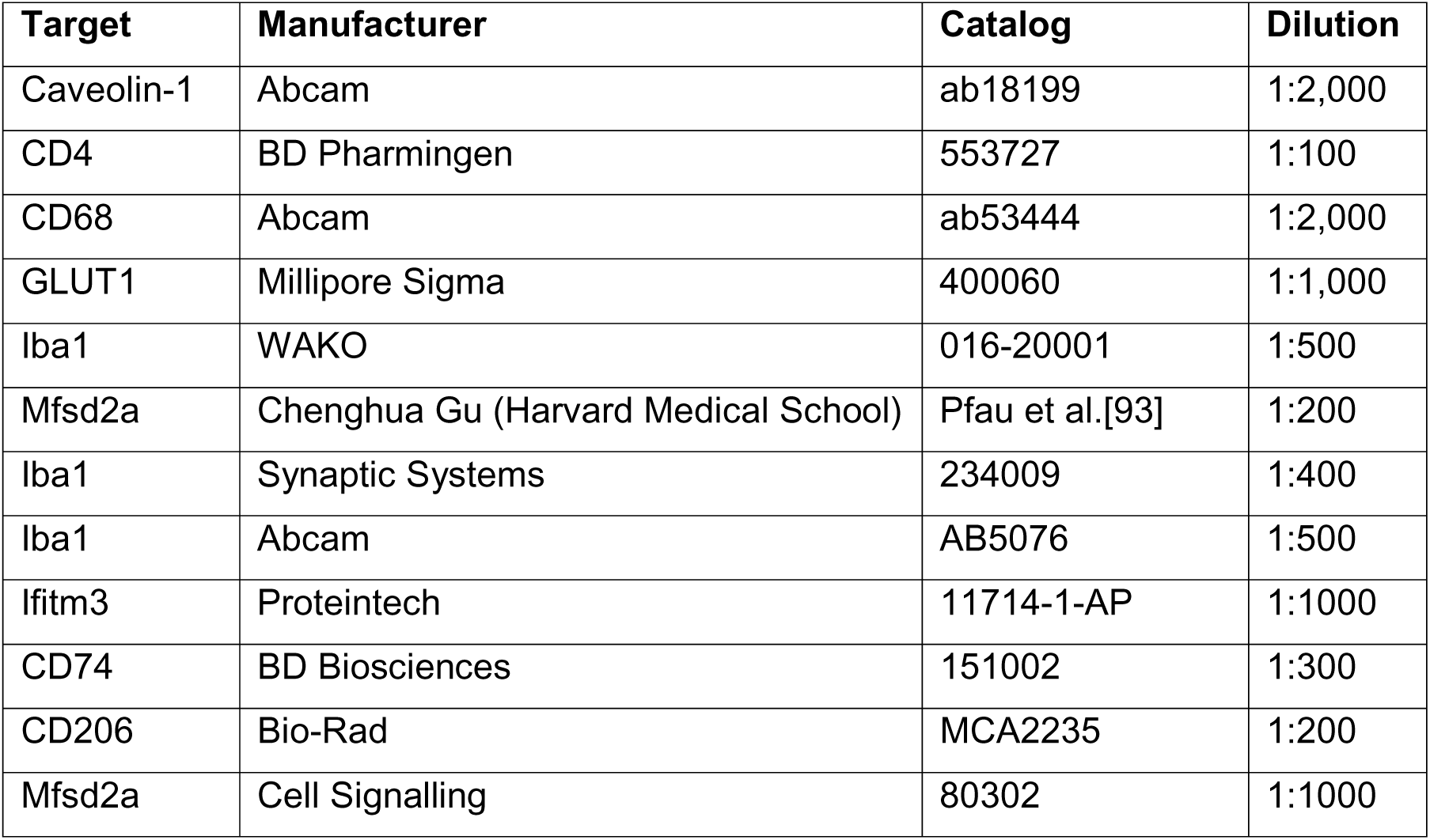
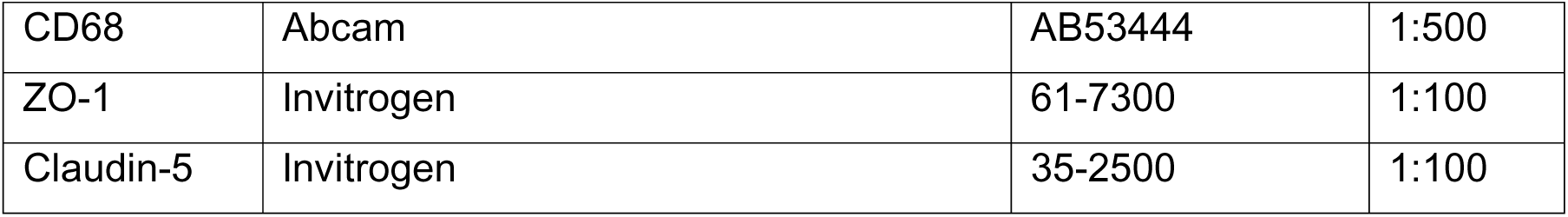

| Target | Manufacturer | Catalog | Dilution |
| --- | --- | --- | --- |
| Caveolin-1 | Abcam | ab18199 | 1:2,000 |
| CD4 | BD Pharmingen | 553727 | 1:100 |
| CD68 | Abcam | ab53444 | 1:2,000 |
| GLUT1 | Millipore Sigma | 400060 | 1:1,000 |
| Iba1 | WAKO | 016-20001 | 1:500 |
| Mfsd2a | Chenghua Gu (Harvard Medical School) | Pfau et al.[93] | 1:200 |
| Iba1 | Synaptic Systems | 234009 | 1:400 |
| Iba1 | Abcam | AB5076 | 1:500 |
| Ifitm3 | Proteintech | 11714-1-AP | 1:1000 |
| CD74 | BD Biosciences | 151002 | 1:300 |
| CD206 | Bio-Rad | MCA2235 | 1:200 |
| Mfsd2a | Cell Signalling | 80302 | 1:1000 |
| CD68 | Abcam | AB53444 | 1:500 |
| ZO-1 | Invitrogen | 61-7300 | 1:100 |
| Claudin-5 | Invitrogen | 35-2500 | 1:100 |

Slides were incubated in primary antibodies at 4°C in a humidified chamber overnight. After three 10-minute washes with PBST, slides were incubated for 2 hours at room temperature in secondary antibodies diluted in PBST with 1% BSA. These were conjugated to AlexaFluor 488 (1:1,000), AlexaFluor 594 (1:1,000), or AlexaFluor 647 (1:500). Following three washes with PBST and two with PBS, slides were cover slipped with Vectashield (Vector Labs, Burlingame, CA) containing the nuclear stain DAPI, then sealed with clear nail polish and stored at -20°C.

### *In situ* hybridization

Plasmids were obtained from Transomic Technologies, and the antisense mRNAs were synthesized using the Digoxigenin RNA Labeling Kit (SP6/T7; Roche, 11175025910). DIG RNA *in situ* hybridization (ISH) and fluorescent *in situ* hybridization (FISH) experiments were performed as described [94, 95]. Mice used for ISH or FISH were perfused for 4 minutes with PBS and brains were dissected out and embedded in O.C.T.

### Multiplexed error-robust *in situ* hybridization (MERFISH)

Mice were anesthetized with isoflurane and intracardially perfused with cold RNAse-free PBS for 4 minutes. Brains were dissected out and immediately embedded in Tissue-Plus O.C.T compound and stored at -80°C until samples could be shipped overnight on dry ice to UMass Chan Medical School. Samples were prepared according to the Vizgen Fresh Frozen Tissue Sample Preparation protocol. Tissue was sectioned in 10 µm slices onto a functionalized coverslip covered with fluorescent beads. Each coverslip contained a section from one PBS sample and a section from one GAS sample. Tissue on coverslips was fixed for 15 minutes at room temperature in 4% paraformaldehyde in PBS, followed by three washes with PBS. Tissue was then permeabilized in 70% ethanol for 24 hours, washed with PBS and incubated with blocking solution for 1 hour, followed by 1 hour of incubation with the primary antibody against vessel marker CD31 (BioLegend, 102502), diluted 1:20 in blocking solution (Vizgen). The tissue was then washed three times with PBS and incubated for 1 hour with an oligo-conjugated secondary antibody diluted 1:1,000 in blocking solution. The sample was washed three times with PBS and fixed for 15 minutes at room temperature in 4% paraformaldehyde in PBS, followed by three washes with PBS. After a 30-minute wash with Formamide Wash Buffer (30% formamide in 2X saline sodium citrate, or SSC) at 37°C, the MERFISH library mix was added and allowed to hybridize for 48 hours. Sample was then washed and incubated at 47°C with Formamide Wash Buffer twice for 30 minutes each and then the tissue was embedded in a polyacrylamide gel followed by incubation with tissue clearing solution (2X SSC, 2% SDS, 0.5% v/v Triton X-100, and proteinase K 1:100) overnight at 37°C. Then, tissue was washed and hybridized for 15 minutes with the first hybridization buffer containing DAPI, polyT and the readout probes associated with the first round of imaging. After washing, the coverslip was assembled into the imaging chamber and placed into the microscope for imaging. MERFISH imaging was performed as previously described[96] with parameter files provided by Vizgen. Briefly, the sample was loaded into a flow chamber connected to the Vizgen Alpha Instrument. First, a low-resolution mosaic image was acquired (405 nm channel) with a low magnification objective (10x). Then the objective was switched to a high magnification objective (60x) and seven 1.5-μm z-stack images of each field of view position were generated in 749 nm, 638 nm and 546 nm channels. A single image of the fiducial beads on the surface of the coverslip was acquired and used as a spatial reference (477 nm channel). After each round of imaging, the readout probes were extinguished, and the sample was hybridized with the next set of readout probes. This process was repeated until combinatorial FISH was completed.

Raw data were decoded using the MERlin pipeline (Vizgen, v0.1.12) using the relevant library codebook. Cell boundaries were segmented in each FOV using a seeded watershed algorithm with DAPI signal as the seed and poly-T signal as the watershed channel. The volume, X position, and Y position of these cell boundaries, as well as the probe counts within each cell boundary, were output for further analysis.

Probes for the following transcripts were used: *Abca7*, *Abcc3*, *Abcg2*, *Ablim1*, *Actb*, *Acvrl1*, *Adam10*, *Adam17*, *Adgrf5*, *Adgrl4*, *Adora1*, *Aff3*, *Ago4*, *Agt*, *Ahr*, *Akap12*, *Aldoc*, *Anxa1*, *Ap2m1*, *Aqp4*, *Arc*, *Arg1*, *Arhgap29*, *Arl15*, *Arpc2*, *Atmin*, *Atp10a*, *Axl*, *Baiap2l1*, *Bard1*, *Bin1*, *Birc5*, *Bmp6*, *Brca1*, *Btk*, *C1qa*, *C1qb*, *C1qbp*, *C1qc*, *C3*, *C3ar1*, *C4a*, *C5ar1*, *Cald1*, *Casp7*, *Casp8*, *Cass4*, *Ccl2*, *Ccl22*, *Ccl3*, *Ccr2*, *Cd14*, *Cd163*, *Cd27*, *Cd33*, *Cd3e*, *Cd4*, *Cd47*, *Cd68*, *Cd72*, *Cd74*, *Cd79a*, *Cd84*, *Cd86*, *Cd8a*, *Cdh5*, *Cdh9*, *Cemip*, *Cenpa*, *Cgnl1*, *Chek2*, *Chit1*, *Cldn5*, *Clec7a*, *Clic4*, *Clu*, *Cmtm8*, *Cobll1*, *Col6a3*, *Cotl1*, *Crim1*, *Csad*, *Csf1r*, *Csf2*, *Csf2ra*, *Csf2rb*, *Cspg4*, *Cstb*, *Ctgf*, *Ctsb*, *Cx3cr1*, *Cxcl10*, *Cyr61*, *Dach1*, *Dapk1*, *Dclre1a*, *Ddx58*, *Des*, *Dlc1*, *Dna2*, *Dock2*, *Dock9*, *Dusp1*, *E2f1*, *Ebf1*, *Ebi3*, *Ece1*, *Edn1*, *Edn3*, *Efnb2*, *Egfl7*, *Egfr*, *Egr1*, *Elovl7*, *Emcn*, *Emp1*, *Eng*, *Enpp6*, *Entpd1*, *Epas1*, *Epb41l4a*, *Epha1*, *Erg*, *Esam*, *Esyt2*, *Fancd2*, *Fbrs*, *Fbxw17*, *Fcer1g*, *Fcgr1*, *Fcgrt*, *Fcrls*, *Fgd2*, *Flcn*, *Flnb*, *Flt1*, *Flt3*, *Flt4*, *Fn1*, *Folr2*, *Fos*, *Foxj1*, *Foxp1*, *Ftsj3*, *Gad1*, *Gad2*, *Galnt18*, *Gbp2*, *Gfer*, *Gna13*, *Gna15*, *Gpi1*, *Gpr183*, *Gpr34*, *Gpr84*, *Grb10*, *Grn*, *Gusb*, *H2-D1*, *Hcar2*, *Heg1*, *Hells*, *Hexb*, *Hmgb2*, *Hmox1*, *Icam1*, *Ifi30*, *Ifih1*, *Igsf6*, *Il1a*, *Il1b*, *Il1rl2*, *Il1rn*, *Il21r*, *Il2rg*, *Il3ra*, *Impact*, *Inpp5d*, *Irak4*, *Irf7*, *Irf8*, *Itga1*, *Itga2*, *Itga6*, *Itgae*, *Itgal*, *Itgam*, *Itgax*, *Itgb1*, *Itgb3*, *Itgb5*, *Itm2a*, *Jcad*, *Jun*, *Kif26a*, *Kit*, *Lacc1*, *Lair1*, *Lama2*, *Lcn2*, *Ldb2*, *Ldlrad3*, *Lef1*, *Lfng*, *Lgals3*, *Lgmn*, *Lig1*, *Liph*, *Lmnb1*, *Lrch3*, *Lrp1*, *Lrrc3*, *Ly6g*, *Ly9*, *Lyn*, *Lyve1*, *Lyz2*, *Mb21d1*, *Mcm2*, *Mcm5*, *Mcm6*, *Mctp1*, *Mecom*, *Mef2c*, *Meg3*, *Mertk*, *Mfsd6l*, *Mgat1*, *Mki67*, *Mkl2*, *Mmp2*, *Mmp9*, *Mpnd*, *Mrc1*, *Ms4a1*, *Ms4a6d*, *Ms4a7*, *Msn*, *Mvp*, *Mybl2*, *Myrip*, *Nampt*, *Napsa*, *Ncaph*, *Ncf1*, *Nckap1l*, *Nebl*, *Nfib*, *Nlrp3*, *Nos3*, *Nostrin*, *Nrm*, *Ocln*, *Olfm2*, *Olfml3*, *Osm*, *Osmr*, *P2ry12*, *Palmd*, *Pam*, *Pard3*, *Pcna*, *Pde10a*, *Pde4d*, *Pdgfra*, *Pdgfrb*, *Pdlim5*, *Pdpn*, *Pdzrn3*, *Pecam1*, *Picalm*, *Pik3cg*, *Pla2g4a*, *Plac8*, *Plcb4*, *Plcg2*, *Plekha6*, *Plekhg1*, *Plp1*, *Plpp1*, *Pltp*, *Plxdc2*, *Plxna2*, *Podxl*, *Pole*, *Ppfibp1*, *Prdx5*, *Prickle2*, *Prkg1*, *Pros1*, *Psmb8*, *Ptk2b*, *Ptpn6*, *Ptprc*, *Ptprg*, *Ptprj*, *Ptprm*, *Pvalb*, *Qars*, *Rad23b*, *Rad51*, *Rae1*, *Rapgef4*, *Rbms3*, *Rngtt*, *Rora*, *Rorb*, *Rrm2*, *Rsad2*, *Rundc3b*, *Sall1*, *Sardh*, *Sash1*, *Sdf4*, *Sell*, *Serpina3n*, *Serpine1*, *Serpinf1*, *Siglecf*, *Slamf8*, *Slamf9*, *Slc17a6*, *Slc1a1*, *Slc39a10*, *Slc40a1*, *Slc4a4*, *Slc7a1*, *Slco2b1*, *Slfn8*, *Smc3*, *Snx2*, *Sorbs1*, *Sorbs2*, *Sox10*, *Sox2*, *Sox9*, *Spi1*, *Spp1*, *Sptbn1*, *Srgn*, *Sst*, *St8sia6*, *Stat1*, *Syk*, *Syne1*, *Syne2*, *Tacc1*, *Tead1*, *Tek*, *Tgfa*, *Tgfbi*, *Tgfbr1*, *Tgfbr2*, *Tgm2*, *Thsd4*, *Timeless*, *Timp2*, *Timp3*, *Tlr2*, *Tlr4*, *Tmem119*, *Tmem173*, *Tmtc1*, *Tnfrsf11a*, *Tnfrsf1a*, *Tnfsf13b*, *Top2a*, *Trem1*, *Trem2*, *Trim47*, *Tshz2*, *Tspan33*, *Ttll12*, *Ttr*, *Txnrd1*, *Tyrobp*, *Unc13b*, *Ung*, *Utrn*, *Vac14*, *Vav1*, *Vcl*, *Vegfa*, *Vim*, *Vip*, *Vtn*, *Vwf*, *Was*, *Wwtr1*, *Xpo1*, *Zbp1*, *Zbtb46*.

### Flow cytometry

Mice were anesthetized with isoflurane and intracardially perfused with PBS for 4 minutes. Brain, as well as combined nasal associated lymphoid tissue/olfactory epithelium (NALT/OE), were dissected out, placed in cold Dulbecco’s Modified Eagle’s Medium (DMEM) (Genesee, 25-500), and pressed through a cell strainer with the end of a sterile syringe. Samples were collected in 10 mL of a 30% Percoll (Cytiva, GE17-0891-01) suspension in DMEM, and underlaid with 1 mL of 70% Percoll. Spleen samples were suspended in 3 mL of Red Blood Cell Lysis Buffer (155 mM NH_4_Cl, 10 mM KHCO_3_, 0.1 mM EDTA) at room temperature for 10 minutes. Samples were centrifuged at 4°C at 1300 rcf for 30 minutes, then immune cells were collected at the interface. All samples were then filtered through a cell strainer, washed with 2 mL DMEM, and centrifuged for 10 minutes at 800 rcf, and resuspended in T cell media (RPMI with fetal bovine serum (FBS) 1:10, penicillin/streptomycin 1:100, MEM-NEAA 1:100, glutamine 1:100, 55 µM β-mercaptoethanol 1:1,000) with 1X Cell Stimulation Cocktail (plus protein transport inhibitors) (eBioscience, 00-4975-93). Samples were incubated at 37°C for 4 hours, washed with FACS buffer, and incubated in anti-CD16/CD32 Fc block (BD Biosciences, 553141, 1:200) for 15 minutes on ice. All subsequent steps were performed at 4°C. After washing with FACS buffer, samples were incubated in cell surface stains (a complete list of antibodies is included in table below) with either Fixable Viability Dye 780 (Invitrogen, 65086518, 1:4,000) or Live/Dead Aqua (ThermoFisher, L34965, 1:1,000) in the dark for 1 hour, fixed for 30 minutes using the Intracellular Fixation & Permeabilization kit (eBioscience, 88-8824-00) according to manufacturer instructions. Intracellular stains were diluted in 1X permeabilization buffer and incubated for 1 hour, then washed in permeabilization buffer and resuspended in FACS buffer.

Samples were analyzed in the Columbia Stem Cell Initiative Flow Cytometry Core. Time course experiments were analyzed using a ZE-5 analyzer (Bio-Rad, Hercules, CA) equipped with 355 nm, 405 nm, 488 nm, 561 nm, and 640 nm lasers. Compensation controls used splenocytes for most cell surface markers and compensation beads (BD Biosciences, 552844 for rat hosts; Thermo Fisher, 01-3333-41 for mouse or Armenian hamster hosts) for cytokines. All other flow cytometry experiments were analyzed using a NovoCyte Penteon (Agilent, Santa Clara, CA) equipped with 349 nm, 405 nm, 488 nm, 561 nm and 637 nm lasers. Compensation controls used splenocytes for live/dead control and compensation beads for all other markers.

All flow cytometry data analysis was performed using FlowJo 10.5 (FlowJo, LLC). Gates for forward and side scatter, singlets, live cells and CD4^+^ T cells were set by eye, while all other gates were set using fluorescence minus one (FMO) controls, with a typical cutoff of <1% of the population.

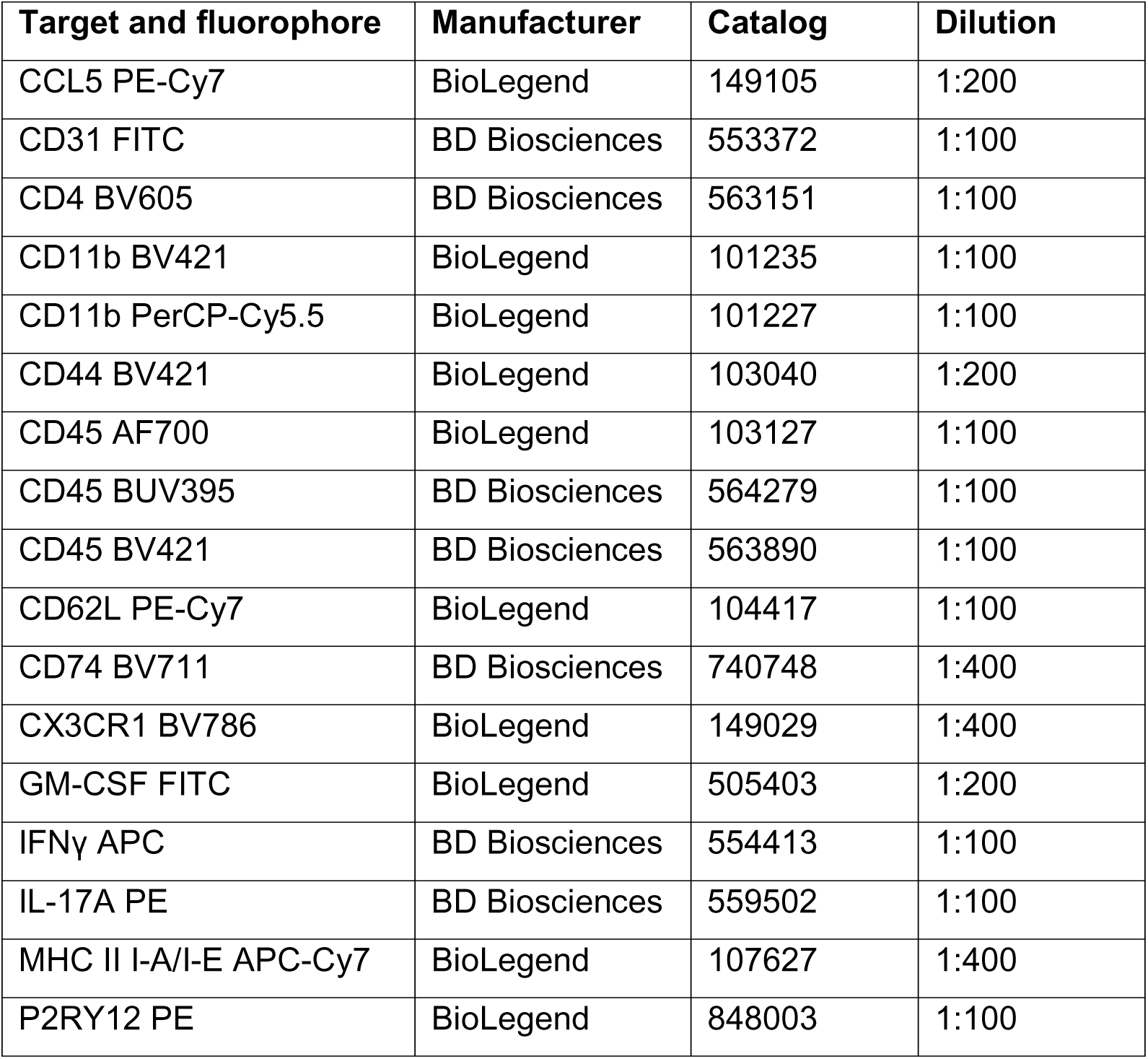

### Cell culture

Primary mouse brain endothelial cells (mBECs; Cell Biologics, C57-6023) and primary human microvascular endothelial cells (HBMEC; Cell System, ACBRI 376) were cultured as monolayers at 37°C with 5% CO_2_ and used to evaluate the effect of cytokines *in vitro*. Cells were grown to confluence on Collagen IV and Fibronectin-coated (Corning, CB-40233, 356008) dishes in endothelial cell media (Cell Biologics, M1168) for mBECs and Endothelial Cell Basal Medium MV2 (PromoCell, C-22221) supplemented with 10% FBS (Cytiva, SH30071.03) and supplements recommended by the supplier for HBMECs. One day prior to cytokine treatment, cells were switched to 2% FBS without growth factor supplements. Media containing either mouse IFNγ (R&D Systems, 485-MI-100), mouse IL-17A (R&D Systems, 7956-ML-025), mouse GM-CSF (R&D Systems, 415-ML-010) or vehicle (PBS with Ca^2+^ and Mg^2+^) was applied to corresponding wells.

Transendothelial electrical resistance (TEER) was measured in real time using an electric cell-substrate impedance sensing (ECIS) instrument (Applied BioPhysics, ZTheta 96 Well Array Station) as previously described [95]. mBECs and HBMECs were plated on 96-well plates containing electrode arrays (Applied BioPhysics, 96W20idf). Cytokines (Il-17A, GM-CSF, IFNγ, CCL2) were added at a concentration of 50 ng/mL, except IL-1β (R&D Systems, 401-ML) and TNF (R&D Systems, 410-MT), which were added at 10ng/mL. Resistance was monitored over 24 hours from the start of cytokine treatment. The area under the curve (AUC) was calculated for each condition and normalized to AUC of vehicle-treated cells, with values for each independent experiment compared by one-way ANOVA.

Albumin transcytosis was measured *in vitro* by culturing mBECs on collagen IV-coated 3.0-µm PET membrane inserts for 24-well plates (Corning, 353096). Cytokines or vehicle were added at a concentration of 100 ng/mL, with 500 ng/mL lipopolysaccharide (LPS) used as a positive control. Cells were switched to endothelial cell media with 2% FBS and without phenol. Media added to wells contained bovine serum albumin at 400 µg/mL, while media added to inside of transwell inserts contained albumin conjugated to Alexa Fluor 647 (Thermo Fisher, A34785) at 400 µg/mL. Cells were incubated at 37°C with flow-through samples collected from bottom wells at 30 minutes, 1, 2, 4 and 6 hours). Absorbance of flow-through was quantified using the accuSkan FC plate reader (Fisher Scientific, 14-377-576). Background fluorescence was subtracted, and AUC normalized to untreated for each experiment.

### Multiplex immunoassays

#### Mouse olfactory bulb multiplex immunoassay

Twenty-four hours after the final GAS inoculation, pairs of OBs were dissected and flash frozen in liquid nitrogen, then stored at -80°C. Samples were pulverized on ice in cell lysis buffer (Abcam, ab152163) with protease inhibitor cocktail (Thermo Fisher, 78440) and EDTA, using an electric pestle. To remove detergents that may interfere with downstream analysis, the resulting supernatant was dialyzed overnight against PBS with a 2 kDa cassette (Thermo Fisher, 66205). After normalization to total protein concentration by Pierce bicinchoninic acid assay (Thermo Fisher, 23225), analytes were measured by the Irving Institute for Clinical and Translational Research Biomarkers Core Laboratory using a custom mouse Luminex panel (Thermo Fisher, PPX-12-MXEPUF3). Samples were run in duplicate, and standard curves were generated for each analyte. Undetectable values were replaced with half of the lower detection limit for purposes of statistical comparison.

#### Patient serum multiplex immunoassay

Serum protein concentrations were measured by the Irving Institute for Clinical and Translational Research Biomarkers Core Laboratory using a 45-Plex Luminex assay (Invitrogen, EPX260-26088-901). Samples were run in duplicate, and standard curves were run for each analyte. Undetectable values were replaced with half of the lower detection limit (**Supplementary Data Table 6**) for statistical comparisons.

### Quantification and statistical analysis Analysis of scRNAseq data

Single-cell RNA sequencing data (**Supplementary Data Table 1**) was analyzed using Seurat package v4.4.1[97] in RStudio. Upon data import, genes detected in fewer than three cells, and cells with fewer than 200 genes were excluded. Cells were removed from the merged data set if they had fewer than 1,000 or more than 50,000 molecules detected, or greater than 20% mitochondrial reads. Data was normalized and highly variable features identified using default parameters, then scaled, followed by linear dimensional reduction using PCA. Dimensionality of the data was selected using the Elbow plot method with 50 dimensions, and cells were clustered with a resolution of 1 for OB, and 0.4 for endothelial cells and microglia. Dimensionality reduction for visualization was performed with uniform manifold approximation and projection (UMAP). The Harmony package v1.2[98] was used for batch correction.

Cluster identity was assigned using the following cell type markers: neurons (*Map2*, *Snap25*), astrocytes (*Gfap*, *Aqp4*), olfactory ensheathing cells (*Frzb*), oligodendrocytes/oligodendrocyte precursor cells (*Pdgfra*), endothelial cells (*Cldn5*, *Pecam1*), pericytes (*Pdgfrb*, *Atp13a5*), fibroblasts (*Col1a1*, *Fbln1*), microglia (*Tmem119*, *P2ry12*), macrophages (*Aif1*, *Plac8*), neutrophils (*Ly6g*, *Camp*), dendritic cells (*Xcr1*, *Ccr9*, *Cd209a*), B cells (*Cd19*), CD4 T cells (*Cd4*), CD8 T cells (*Cd8a*), NK cells (*Klrb1c*), and B cells (*Cd19*).

Differential expression analysis was performed using a mixed-effects model algorithm (MAST) [99] to avoid pseudo-replication bias [100]. Expression from biological replicates was aggregated for heatmap visualization and significance by mixed-effects analysis displayed by formatting of gene names. Expression of *ex vivo* activation genes (*Dusp1*, *Fos*, *Hist1h1d*, *Hist1h2ac*, *Jun*, *Nfkbid*, *Nfkbiz*) were added using AddModuleScore and plotted against *Ccl3* and *Ccl4* using FeatureScatter. Additional analysis was performed with BB Browser 3 (BioTuring) software. Signature scores were plotted in BBrowser3 using the following pathway markers: Antigen presentation (*B2m*, *Cd74*, *H2-Aa*, *H2-Ab1*, *H2-D1*, *H2-Eb1*, *H2-K1*, *H2-Q4*, *H2-Q6*, *H2-Q7*, *Tap1*, *Tap2*), disease-associated microglia (*Apoe*, *Axl*, *Cd9*, *Csf1*, *Cst7*, *Itgax*, *Lpl*, *Spp1*, *Tyrobp*), homeostatic microglia (*Cd33*, *Cst3*, *Cx3cr1*, *Fcrls*, *Gpr34*, *Olfml3*, *P2ry12*, *P2ry13*, *Sall1*, *Tmem119*), and interferon signaling (*Ifi30*, *Ifi204*, *Ifi211*, *Ifit1*, *Ifitm3*, *Irf1*, *Irf7*, *Isg15*, *Oas1a*, *Stat1*, *Stat2*).

Gene set enrichment analysis (GSEA)[37, 38] with curated pathways was performed using was performed using curated and database-derived gene lists for blood-brain barrier, response to LPS, inflammation, extracellular matrix, interferon response, antigen presentation, chemokine and cytokine signaling, endothelial cell proliferation, endothelial cell migration, disease-associated microglia, apoptosis, leukocyte chemotaxis and phagocytosis (**Supplementary Data Table 4**). Analysis was run using the GSEA desktop tool (Broad Institute, v4.1.0), using pre-ranked weighted settings. GSEA using gene ontology (GO) pathway lists was performed using the enrichplot R package v1.24.4 (**Supplementary Data Table 5**). Cross entropy analysis was performed on UMAP coordinates using the Cross-Entropy-test [34] in R. Receptor-ligand analysis was performed using CellChat v1.6.1 [51, 52].

### Analysis of MERFISH data

MERFISH data was analyzed in RStudio using Seurat 4.1.0.9005, R 4.0.0 and custom-made scripts as previously described [101]. Cell segmentations with volume < 50µm^3^ or < 10 unique transcripts were first excluded. Cell gene expression data of each cell was then normalized to that cell’s volume and the total transcript count of that cell, then scaled. To correct for global differences in total transcript counts between coverslips (each containing one GAS sample and one PBS sample), we performed ComBat [102] batch correction (sva 3.38.0).

To identify individual cell types, we performed principal component analysis was performed using the entire probe library (391 transcripts) as the variable features, followed by linear dimensional reduction. Dimensionality of the data was selected using the jackstraw method with 28 dimensions, and cells were clustered with a resolution of 2.4. Dimensionality reduction for visualization was performed with uniform manifold approximation and projection (UMAP). Clusters were manually annotated based on the spatial distribution of the cells in the tissue and the expression cell type-specific marker genes: neurons (*Meg3, Gad1*), astrocytes (*Aqp4, Sox9*), olfactory ensheathing cells (*Plp1, Cldn5*), oligodendrocytes/oligodendrocyte precursor cells (*Pdgfra, Sox10*), endothelial cells (*Cldn5*, *Itm2a*), pericytes (*Pdgfrb*), fibroblasts (*Cemip*), microglia (*Tmem119*, *P2ry12*), macrophages (*Mrc1*), neutrophils (*Itgal*, *Mmp9*), and T cells (*Cd3e*, *Cd4, Cd8a*).

Because of imperfections in cell boundary segmentation, a small fraction of cells expressed cell type markers for multiple cell types. Raw images of a subset of these cells were visually inspected using MERSCOPE Visualizer software (Vizgen, 2.1.2589.1) to confirm that these clusters were due to cell segmentation errors (typified by two distinct clusters of cell-type specific transcripts within the same cell boundary). Clusters composed of these “hybrid” cells were removed from the analysis, and embedding and clustering analysis were iteratively repeated until all “hybrid” clusters were removed.

The glomerular, external plexiform, and granular layers of the OB for each sample were outlined using MobileFish and coordinates recorded for point-in-polygon analysis and regional assignment of microglia. Raw counts were normalized to the PBS condition for each batch and used for gene expression analysis. Endothelial cell gene expression was compared on log_2_ fold change values using a one-sample t test. Microglia gene expression between the glomerular and granular layers used a ratio t test. Nearest neighbor analysis of microglial distance to T cells was calculated based the x,y coordinates of the centers of the cell segmentations using a custom python script.

### Statistical analysis

Most statistical analyses were performed by GraphPad Prism 10.3.0. All tests were two-sided using a significance level α = 0.05. Outliers were identified and excluded using the ROUT method (Q = 1%). Significance was notated as ns, p > 0.05; *, p < 0.05; **, p < 0.01; ***, p < 0.001. Error bars represent mean with SEM throughout.

### Immunofluorescence quantification

#### Quantification of microglial number

Three OB sections, corresponding to bregmas 4.5, 4.28 and 3.92, were imaged using a Zeiss AxioImager microscope at magnification 20x for each animal. The number of Iba1^+^CD68^+^ cells (corresponding to activated microglia) in the glomerular layer was manually counted, and averaged across the three sections. Iba1^+^ cells (microglia) were considered CD68^+^ if the CD68 fluorescence occupied more than 50% of the cell surface area, as previously described[23, 25].

#### Quantification of BBB leakage

Bregma 4.28 sections were imaged using a Zeiss AxioImager microscope at magnification 10x. Using ImageJ [103], 20 small, rectangular regions of interest (ROIs) were placed around the glomerular layer of the OB, avoiding the vasculature, and average fluorescence quantified for each animal. The same process was repeated for the granular layer.

#### Quantification of BEC an microglial marker expression

Tiled images of bregma 4.28 sections were taken at 20x using a Zeiss LSM700 or LSM900 confocal microscope and Zeiss AxioImager M2, and maximum intensity projections were created if it is possible. Using FIJI, ROIs across the whole OB were selected using Otsu thresholding on a vessel marker. Then average fluorescence intensity was measured within the ROIs. Due to neuronal expression of *Itih5* in the glomerular layer, ROIs were restricted to the granular layer of the OB for quantification. For the microglial marker analyses, Iba1 positive cells were used for generating a mask and CD74, CD68, Ifitm3 were measured within Iba1^+^ microglial cell mask.

## Data availability

Raw sequencing data, metadata and count tables for scRNAseq samples have been made available in the Gene Expression Omnibus (GEO) under GSE221724 and GSE221106. The MERFISH raw output files were not deposited in a public repository due to storage restrictions but can be made available upon request.

## Code availability

The code to reproduce this study’s processed data is available at GitHub.

