## Supplementary Information for "Th17 effector cytokines induce shared and distinct microglial and endothelial cell responses in post-streptococcal encephalitis"

[1]Department of Neurology, Columbia University Irving Medical Center, New York, NY, 10032, USA
[2]Department of Neurobiology, Brudnick Neuropsychiatric Research Institute, University of Massachusetts Chan Medical School, Worcester, MA, 01605, USA
[3]Department of Neurology, Thomas Jefferson University, Philadelphia, PA, 19107, USA
[4]Lyme and Tick-Borne Diseases Research Center, Department of Psychiatry, Columbia University Irving Medical Center, New York, NY, 10032, USA
[5]Section of Behavioral Pediatrics, National Institute of Mental Health, Bethesda, MD, 20814, USA
[6]Department of Pathology and Cell Biology, Columbia University Irving Medical Center, New York, NY, 10032, USA
[7]These first authors contributed equally to the manuscript.
[8]These senior authors jointly supervised the manuscript.

### Supplementary Information

#### Extended Data Figures and Figure Legends

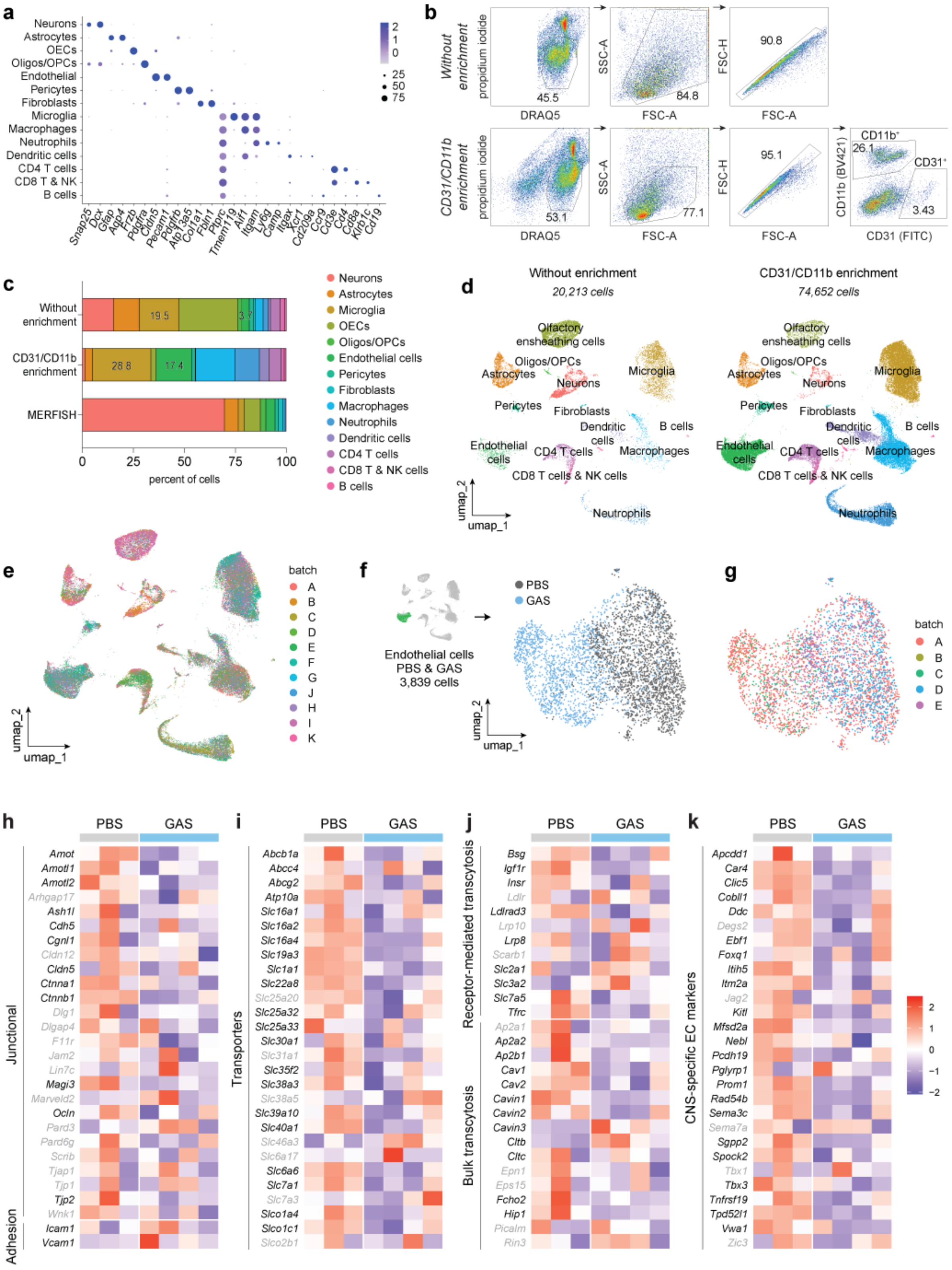

**Extended Data Figure 1. Identification of olfactory bulb cell types isolated from PBS and GAS-infected mice for scRNAseq, and transcriptional shifts in brain endothelial cells after multiple intranasal GAS infections.**

**a,** Dot plot showing canonical molecular markers used to assign cell identities in the olfactory bulb (OB) from PBS- and Group A Streptococcus (GAS)–infected mice for single-cell RNA sequencing (scRNA-seq). Color intensity indicates mean expression, and dot size denotes the percentage of cells expressing each gene. **b,** Fluorescence-activated cell sorting (FACS) gating strategy for isolation of myeloid cells and brain endothelial cells (BECs) from the OB. Top panels show ungated samples; bottom panels show enrichment for $CD31^+$ BECs and $CD11b^+$ myeloid cells. **c,** Cell type composition of OB samples used for scRNA-seq (with and without enrichment) and MERFISH analyses. Numbers within bars indicate the fraction of BECs (green) and microglia (gold) among all isolated cells. **d,** UMAP visualization of scRNA-seq data from OBs of PBS- and GAS-infected mice without enrichment (left) and following enrichment for $CD31^+$ BECs and $CD11b^+$ myeloid cells (right). **e,** UMAP plot showing all identified cell clusters across samples, colored by sequencing batch.**f, g,** UMAP plots of BECs colored by experimental condition (PBS, gray; GAS, light blue) (**f**) and by sequencing batch (**g**).**h–k,** Heat maps of differentially expressed genes in BECs from PBS (gray) and GAS-infected (light blue) mice, grouped by functional category: blood–brain barrier junction and adhesion proteins (**h**), transporters (**i**), transcytosis-related genes (**j**), and brain endothelial identity markers (**k**). Values are shown as log(z-scores); red indicates higher and blue lower relative expression. Genes labeled in black are significant (adjusted $P < 0.05$); gray labels indicate non-significant genes.

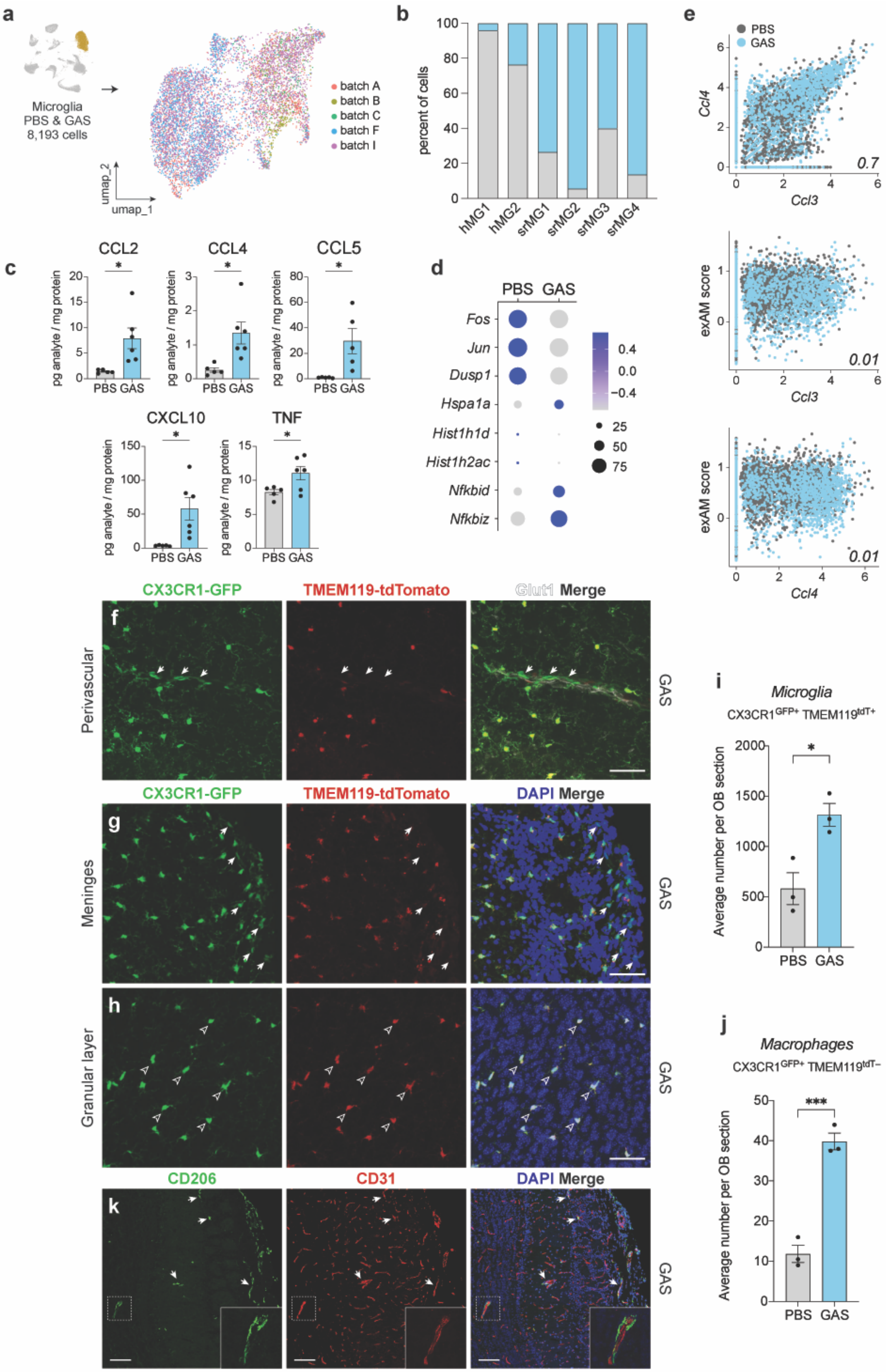
a
Microglia
PBS & GAS
8,193 cells
batch A
batch B
batch C
batch F
batch I
umap_1
umap_2
b
percent of cells
hMG1
hMG2
srMG1
srMG2
srMG3
srMG4
e
PBS
GAS
Ccl4
Ccl3
0.7
exAM score
0.01
c
CCL2
CCL4
CCL5
CXCL10
TNF
pg analyte / mg protein
PBS GAS
d
Fos
Jun
Dusp1
Hspa1a
Hist1h1d
Hist1h2ac
Nfkbid
Nfkbiz
CX3CR1-GFP
TMEM119-tdTomato
Glut1 Merge
DAPI Merge
Perivascular
Meninges
Granular layer
GAS
CD206
CD31
i
Microglia
CX3CR1GFP+ TMEM119tdT+
Average number per OB section
j
Macrophages
CX3CR1GFP+ TMEM119tdT−

**Extended Data Figure 2. Microglia upregulate disease-associated and chemokine genes in the olfactory bulb following repeated intranasal GAS infection.**

**a,** UMAP plot showing sequencing batch identity of microglia isolated from olfactory bulbs (OBs) of PBS- and Group A Streptococcus (GAS)–infected mice. **b,** Stacked bar plot indicating the proportion of microglia derived from PBS (gray) and GAS-infected (light blue) mice within each identified microglial subpopulation. **c,** Dot plots showing concentrations of selected cytokines (pg/mg tissue) measured in whole OB lysates by multiplex immunoassay. Comparisons between PBS and GAS-infected mice were performed using an unpaired *t*-test with Welch's correction ($P < 0.05$; $n = 5$ PBS and $n = 6$ GAS). Data are shown as mean ± SEM. **d,** Dot plot showing expression of ex vivo activation (exAM) gene signatures in microglia isolated from PBS- and GAS-infected OBs. Dot size represents the percentage of microglia expressing each gene, and color intensity indicates mean expression level. **e,** Scatter plots of *Ccl3* and *Ccl4* mRNA expression (x-axis) versus exAM module score (y-axis) in individual microglia. Pearson correlation coefficients are shown in the lower right corner of each plot. **f–h,** Representative immunofluorescence images of perivascular macrophages (**f**), meningeal macrophages (**g**), and microglia (**h**) in OBs from GAS-infected $CX3CR1^{GFP}$/ $TMEM119^{tdTomato}$ reporter mice. White arrows (**f, g**) indicate macrophages ($CX3CR1^{GFP}$+ $TMEM119^{tdTomato}$-), and arrowheads (**h**) indicate microglia ($CX3CR1^{GFP}$+ $TMEM119^{tdTomato}$+). Scale bars, 50 μm. **i, j,** Quantification of the average number of microglia (**i**) and macrophages (**j**) in OB sections at bregma positions 4.5, 4.28, and 3.92 from PBS- and GAS-infected $CX3CR1^{GFP}$/ $TMEM119^{tdTomato}$ reporter mice ($n = 3$ mice per group). Statistical comparisons were performed using an unpaired *t*-test ($P < 0.05$; $^{**}P < 0.001$). Data are shown as mean ± SEM. **k,** Representative immunofluorescence images of $CD206^{+}$ perivascular macrophages in OBs from GAS-infected mice. CD206 marks perivascular macrophages (white arrows), and CD31 labels blood vessels. Insets show higher-magnification views of $CD206^{+}$ macrophages associated with $CD31^{+}$ vasculature. Scale bars, 50 μm.

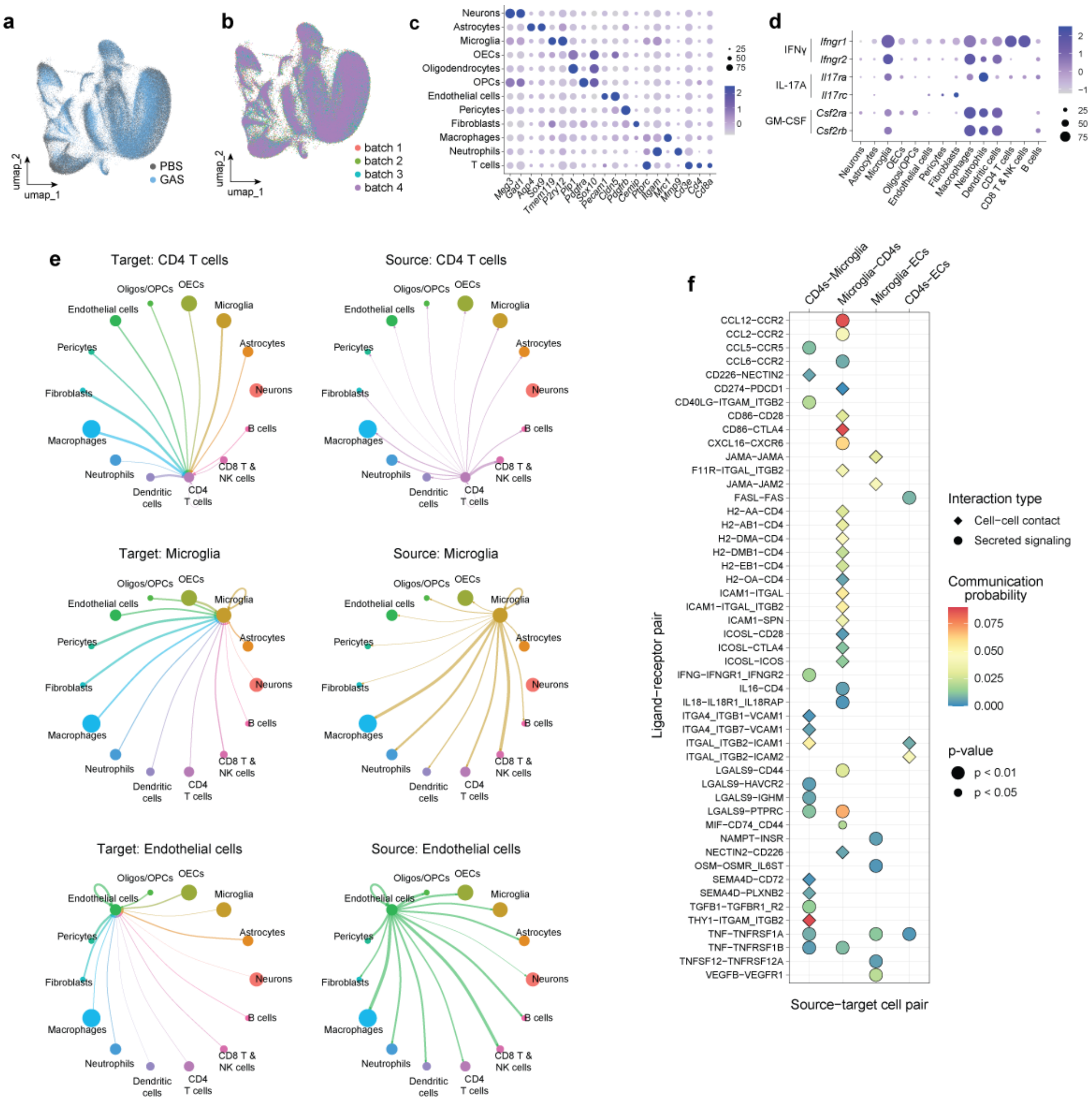


**Extended Data Figure 3. Altered intercellular communication in the olfactory bulb following intranasal GAS infection.**

**a,b,** UMAP visualizations of olfactory bulb (OB) cells profiled by MERFISH, colored by experimental condition (PBS, gray; GAS, blue) (**a**) and by experimental batch (**b**). **c,** Dot plot showing canonical molecular markers used to assign OB cell identities in MERFISH experiments. **d,** Dot plot showing cytokine receptor expression across OB cell types based on scRNA-seq data following intranasal GAS infection. In **c** and **d**, color intensity indicates mean expression and dot size denotes the percentage of cells expressing each gene. **e,** Bubble plots depicting inferred receptor–ligand communication networks following intranasal GAS infection, illustrating signaling from CD4$^+$ T cells, microglia, and brain endothelial cells (BECs) to other OB cell types and reciprocal interactions. **f,** Statistically significant inferred receptor–ligand interactions among CD4$^+$ T cells, microglia, and BECs in GAS-infected mice. Secreted signaling interactions are shown as circles and cell–cell contact–dependent interactions as diamonds. Communication probability is represented by color intensity, and statistical significance (*P* value) by symbol size.

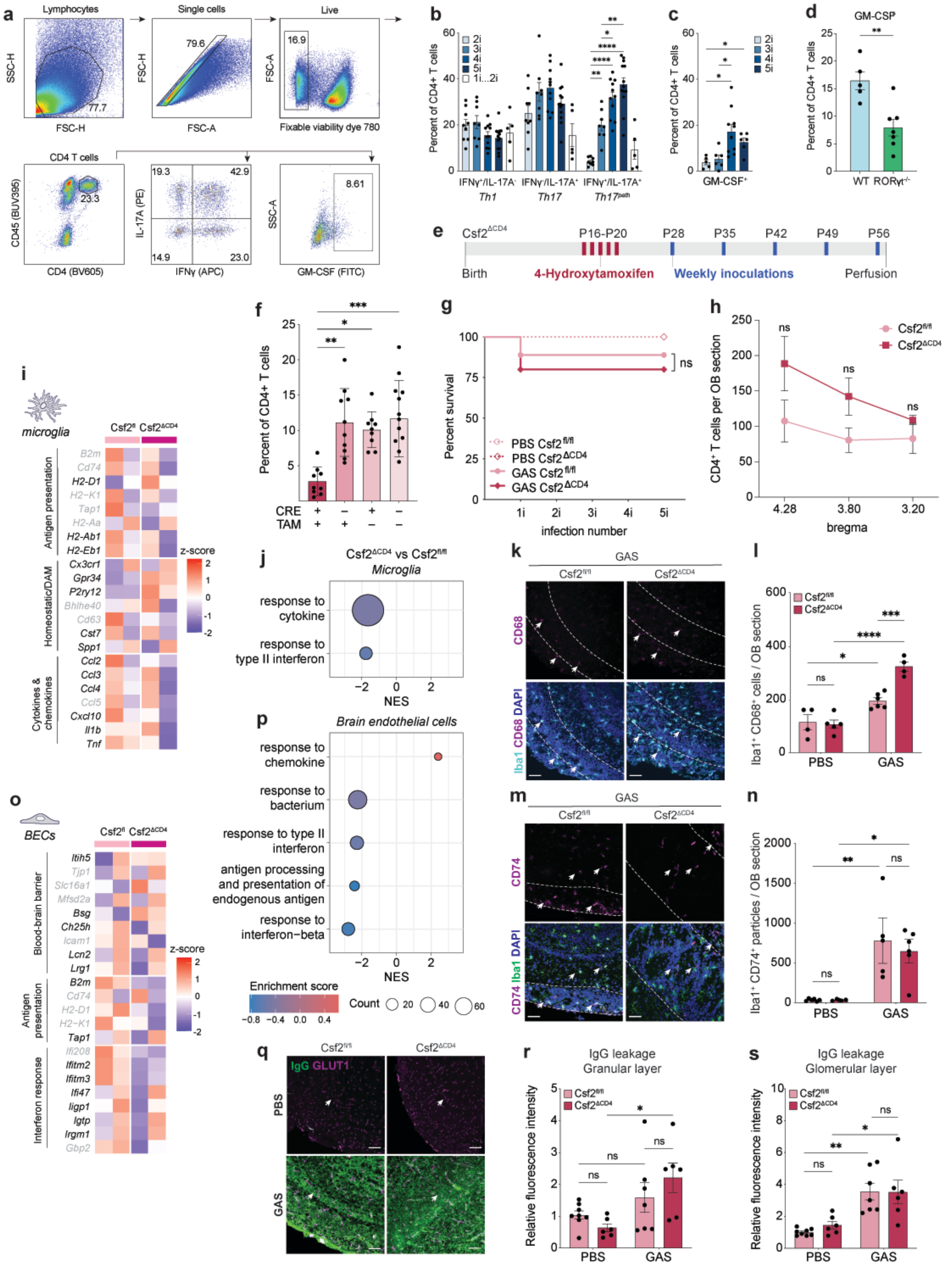

a
Lymphocytes
SSC-H
FSC-H
77.7
Single cells
FSC-H
FSC-A
79.6
Live
FSC-A
Fixable viability dye 780
16.9
CD4 T cells
CD45 (BUV395)
CD4 (BV605)
23.3
IL-17A (PE)
IFNγ (APC)
19.3
42.9
14.9
23.0
SSC-A
GM-CSF (FITC)
8.61
b
Percent of CD4+ T cells
2i
3i
4i
5i
1i...2i
IFNγ⁺/IL-17A⁻
Th1
IFNγ⁻/IL-17A⁺
Th17
IFNγ⁺/IL-17A⁺
Th17path
c
Percent of CD4+ T cells
GM-CSF⁺
d
GM-CSF
Percent of CD4+ T cells
WT
RORγt-/-
e
Csf2ΔCD4
P16-P20
P28
P35
P42
P49
P56
Birth
4-Hydroxytamoxifen
Weekly inoculations
Perfusion
f
Percent of CD4+ T cells
CRE
TAM
g
Percent survival
PBS Csf2fl/fl
PBS Csf2ΔCD4
GAS Csf2fl/fl
GAS Csf2ΔCD4
ns
infection number
h
CD4⁺ T cells per OB section
Csf2fl/fl
Csf2ΔCD4
bregma
4.28
3.80
3.20
i
microglia
Csf2fl
Csf2ΔCD4
Antigen presentation
Homeostatic/DAM
Cytokines & chemokines
B2m
Cd74
H2-D1
H2-K1
Tap1
H2-Aa
H2-Ab1
H2-Eb1
Cx3cr1
Gpr34
P2ry12
Bhlhe40
Cd63
Cst7
Spp1
Ccl2
Ccl3
Ccl4
Ccl5
Cxcl10
Il1b
Tnf
z-score
j
Csf2ΔCD4 vs Csf2fl/fl
Microglia
response to cytokine
response to type II interferon
NES
k
GAS
Csf2fl/fl
Csf2ΔCD4
CD68
Iba1 CD68 DAPI
l
Csf2fl/fl
Csf2ΔCD4
Iba1⁺ CD68⁺ cells / OB section
PBS
GAS
m
GAS
Csf2fl/fl
Csf2ΔCD4
CD74
CD74 Iba1 DAPI
n
Iba1⁺ CD74⁺ particles / OB section
PBS
GAS
o
BECs
Csf2fl
Csf2ΔCD4
Blood-brain barrier
Antigen presentation
Interferon response
Itih5
Tjp1
Slc16a1
Mfsd2a
Bsg
Ch25h
Icam1
Lcn2
Lrg1
B2m
Cd74
H2-D1
H2-K1
Tap1
Ifi208
Ifitm2
Ifitm3
Ifi47
Iigp1
Igtp
Irgm1
Gbp2
z-score
p
Brain endothelial cells
response to chemokine
response to bacterium
response to type II interferon
antigen processing and presentation of endogenous antigen
response to interferon−beta
NES
Enrichment score
Count
q
Csf2fl/fl
Csf2ΔCD4
IgG GLUT1
PBS
GAS
r
IgG leakage
Granular layer
Relative fluorescence intensity
Csf2fl/fl
Csf2ΔCD4
PBS
GAS
s
IgG leakage
Glomerular layer
Relative fluorescence intensity
Csf2fl/fl
Csf2ΔCD4
PBS
GAS


100

**Extended Data Figure 4. $CD4^+$ T cell–derived GM-CSF is required for microglial transcriptional remodeling following repeated intranasal GAS infection.**

**a,** Representative flow cytometry gating strategy for $CD4^+$ T cells isolated from the brain. **b, c,** Dot plots showing the proportion of $IFN\gamma^+IL\text{-}17A^-$ (Th1), $IFN\gamma^-IL\text{-}17A^+$ (Th17), $IFN\gamma^+IL\text{-}17A^+$ ($Th17^{path}$) $CD4^+$ T cells (**b**), and $GM\text{-}CSF^+$ $CD4^+$ T cells (**c**) in the brain after 2–5 intranasal GAS infections. Cells were gated on live singlet $CD45^+CD4^+$ cells. Th1, Th17, and $Th17^{path}$ populations increased with the number of infections. Statistical analysis was performed using one-way ANOVA with Dunnett's T3 multiple-comparisons test (ns, $P > 0.05$; $P < 0.05$; **$P < 0.001$; ***$P < 0.0001$; $n = 9$–12 mice per group). **d,** Quantification of $GM\text{-}CSF^+$ $CD4^+$ T cells in olfactory bulbs (OBs) of GAS-infected wild-type (blue; $n = 5$) and $ROR\gamma t^{-/-}$ (green; $n = 7$) mice. Statistical significance was assessed by unpaired *t*-test (*$P < 0.01$). **e,** Experimental timeline showing 4-hydroxytamoxifen (4-OHT) administration and intranasal GAS infections in $Csf2^{fl/fl}$ and $Csf2^{\Delta CD4}$ mice. **f,** Flow cytometric confirmation of CD4-specific GM-CSF deletion. Nose-associated lymphoid tissue (NALT) from $Csf2^{\Delta CD4}$ ($Cre^+$) mice treated with 4-OHT ($TAM^+$) prior to intranasal GAS infection showed reduced frequencies of $GM\text{-}CSF^+$ $CD4^+$ T cells. Cells were gated on live singlet $CD45^+CD4^+$ cells. Statistical analysis was performed using one-way ANOVA with Sidák's multiple-comparisons test (ns, $P > 0.05$; $P < 0.05$; **$P < 0.001$; ***$P < 0.0001$; $n = 9$–12 mice per group). **g,** Survival curves of PBS- or GAS-infected $Csf2^{fl/fl}$ and $Csf2^{\Delta CD4}$ mice ($n = 9$ mice per group). No significant differences were observed (Mantel–Cox test). **h,** Quantification of $CD4^+$ T cell numbers in the OB at bregma positions 4.28, 3.8, and 3.2 in PBS- and GAS-infected $Csf2^{fl/fl}$ and $Csf2^{\Delta CD4}$ mice. Statistical analysis was performed using mixed-effects analysis with Sidák's multiple-comparisons test (ns, $P > 0.05$; $P < 0.05$; $n = 6$ mice per group). **i, j,** Heat map of differentially expressed microglial genes (**i**) and Gene Ontology (GO) pathway analysis (**j**) comparing GAS-infected $Csf2^{\Delta CD4}$ (maroon) and $Csf2^{fl/fl}$ (pink) mice. **k, l,** Representative immunofluorescence (IF) images and quantification of $Iba1^+CD68^+$ microglia in the glomerular layer of the OB (dashed outlines) from GAS-infected $Csf2^{fl/fl}$ and $Csf2^{\Delta CD4}$ mice. Arrows indicate double-positive cells. **m, n,** Representative IF images and quantification of $Iba1^+CD74^+$ microglia in the glomerular layer of the OB from GAS-infected $Csf2^{fl/fl}$ and $Csf2^{\Delta CD4}$ mice. Arrows indicate double-positive cells. **o, p,** Heat map of differentially expressed brain endothelial cell genes (**o**) and GO pathway analysis (**p**) comparing GAS-infected $Csf2^{\Delta CD4}$ and $Csf2^{fl/fl}$ mice. **q,** Representative IF images showing serum IgG (green) and $Glut1^+$ blood vessels (magenta) in the OB. **r, s,** Quantification of serum IgG leakage in the granular (**r**) and glomerular (**s**) layers of the OB in PBS- and GAS-infected $Csf2^{fl/fl}$ and $Csf2^{\Delta CD4}$ mice. For **i, j, o, p**, values are shown as log(z-scores); red indicates higher and blue lower relative expression. Genes labeled in black are significant (adjusted $P < 0.05$), whereas gray labels indicate non-significant genes. Bars in **l, n, r, s** represent mean ± SEM. Statistical analyses were performed using two-way ANOVA with Šídák's multiple-comparisons test (ns, $P > 0.05$; *$P < 0.01$; **$P < 0.001$; ***$P < 0.0001$; $n$ = 4–8 mice per group).

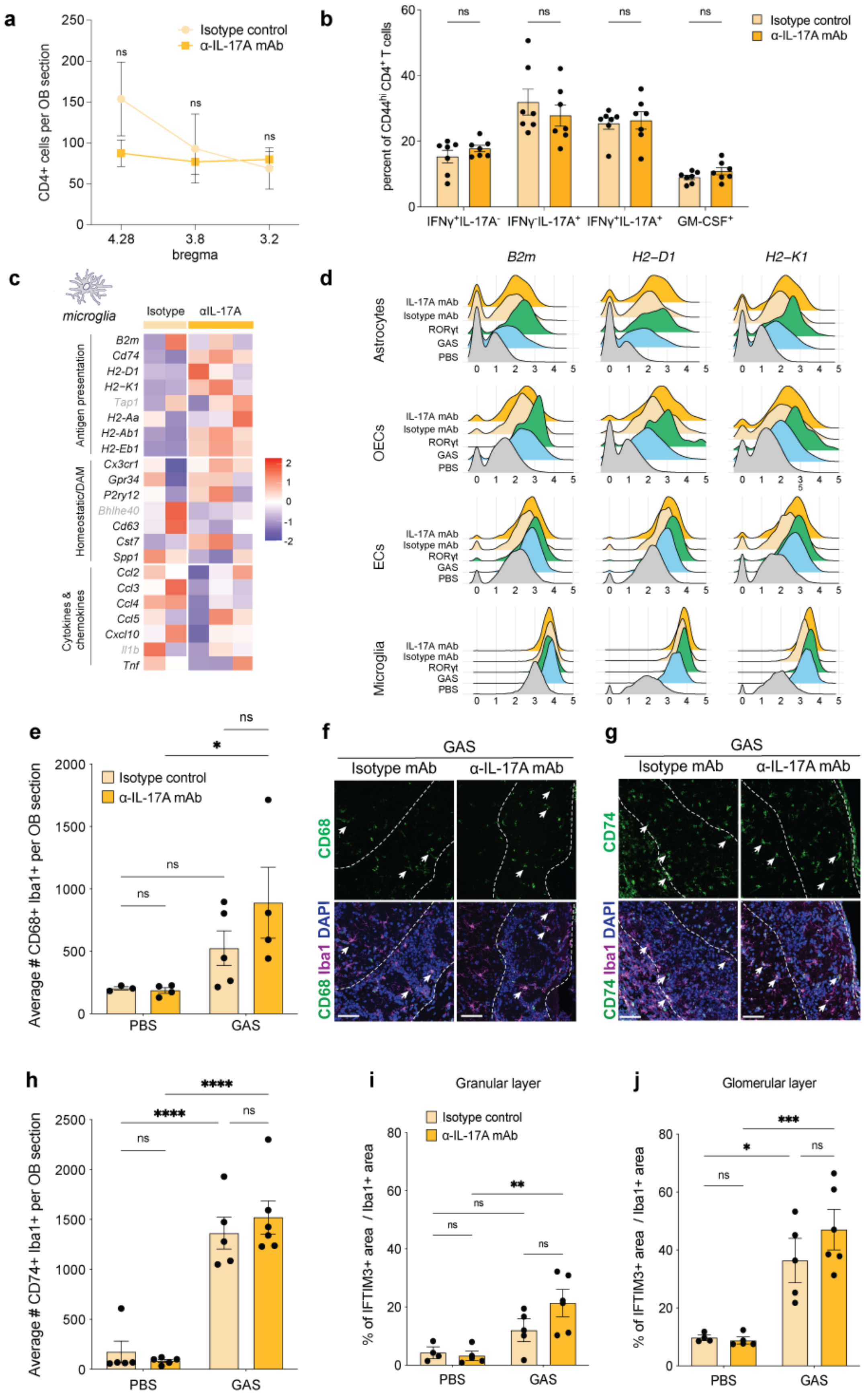

a
Isotype control
α-IL-17A mAb
CD4+ cells per OB section
bregma
4.28
3.8
3.2
ns
b
percent of CD44hi CD4+ T cells
IFNγ+IL-17A-
IFNγ-IL-17A+
IFNγ+IL-17A+
GM-CSF+
c
microglia
Isotype
αIL-17A
Antigen presentation
B2m
Cd74
H2-D1
H2-K1
Tap1
H2-Aa
H2-Ab1
H2-Eb1
Homeostatic/DAM
Cx3cr1
Gpr34
P2ry12
Bhlhe40
Cd63
Cst7
Spp1
Cytokines & chemokines
Ccl2
Ccl3
Ccl4
Ccl5
Cxcl10
Il1b
Tnf
d
B2m
H2-D1
H2-K1
Astrocytes
OECs
ECs
Microglia
IL-17A mAb
Isotype mAb
RORγt
GAS
PBS
e
Average # CD68+ Iba1+ per OB section
Isotype control
α-IL-17A mAb
PBS
GAS
f
GAS
Isotype mAb
α-IL-17A mAb
CD68
CD68 Iba1 DAPI
g
GAS
Isotype mAb
α-IL-17A mAb
CD74
CD74 Iba1 DAPI
h
Average # CD74+ Iba1+ per OB section
PBS
GAS
i
Granular layer
Isotype control
α-IL-17A mAb
% of IFTIM3+ area / Iba1+ area
PBS
GAS
j
Glomerular layer
% of IFTIM3+ area / Iba1+ area
PBS
GAS

**Extended Data Figure 5. IL-17A blockade attenuates microglial antigen presentation and inflammatory gene programs without altering $CD4^+$ T cell infiltration in the olfactory bulb during GAS infection.**

**a,** Quantification of $CD4^+$ T cell numbers in olfactory bulbs (OBs) at bregma positions 4.28, 3.8, and 3.2 from GAS-infected mice treated with isotype control antibody (yellow) or anti–IL-17A monoclonal antibody (α–IL-17A mAb; orange). Statistical analysis was performed using mixed-effects analysis with Sidák's multiple-comparisons test (ns, $P > 0.05$; $n = 4$ isotype control and $n = 6$ α–IL-17A–treated mice). **b, c,** Dot plots showing the proportions of GM-$CSF^+$ $CD4^+$ T cells (**b**) and $CD4^+$ T cell subsets, including Th1, Th17, $Th17^{path}$, and GM-$CSF^+$ $CD4^+$ T cells (**c**), isolated from OBs of GAS-infected mice treated with isotype control (yellow; $n = 7$) or α–IL-17A mAb (orange; $n = 7$). Statistical comparisons were performed using two-way ANOVA with Šídák's multiple-comparisons test. **d, e,** Heat map of differentially expressed microglial genes (**d**) and Gene Ontology (GO) pathway analysis (**e**) comparing GAS-infected mice treated with isotype control (yellow) or α–IL-17A mAb (orange). **f,** Ridge plots showing expression of selected major histocompatibility complex (MHC) class I antigen-presentation genes across OB cell types in wild-type PBS-treated (gray), wild-type GAS-infected (blue), *RORγt*$^{-/-}$ GAS-infected (green), isotype control antibody–treated GAS-infected (yellow), and α–IL-17A mAb–treated GAS-infected (orange) mice. **g–j,** Representative immunofluorescence images and quantification of $Iba1^+CD68^+$ and $Iba1^+CD74^+$ microglia in the glomerular layer of the OB (dashed outlines) from GAS-infected mice treated with isotype control (yellow) or α–IL-17A mAb (orange). Arrows indicate double-positive cells. **k, l,** Quantification of $Ifitm3^+Iba1^+$ microglia in the granular (**k**) and glomerular (**l**) layers of the OB from PBS- or GAS-infected mice treated with isotype control (yellow) or α–IL-17A mAb (orange). For **d** and **e**, values are shown as log(z-scores); red indicates higher and blue lower relative expression. Genes labeled in black are significant (adjusted $P < 0.05$), whereas gray labels indicate non-significant genes. Graph bars represent mean ± SEM. Statistical analyses for all panels were performed using two-way ANOVA with Šídák's multiple-comparisons test (ns, $P > 0.05$; $P < 0.05$; $^*P < 0.01$; $^{**}P < 0.001$; $^{***}P < 0.0001$; $n = 3$–6 mice per group).

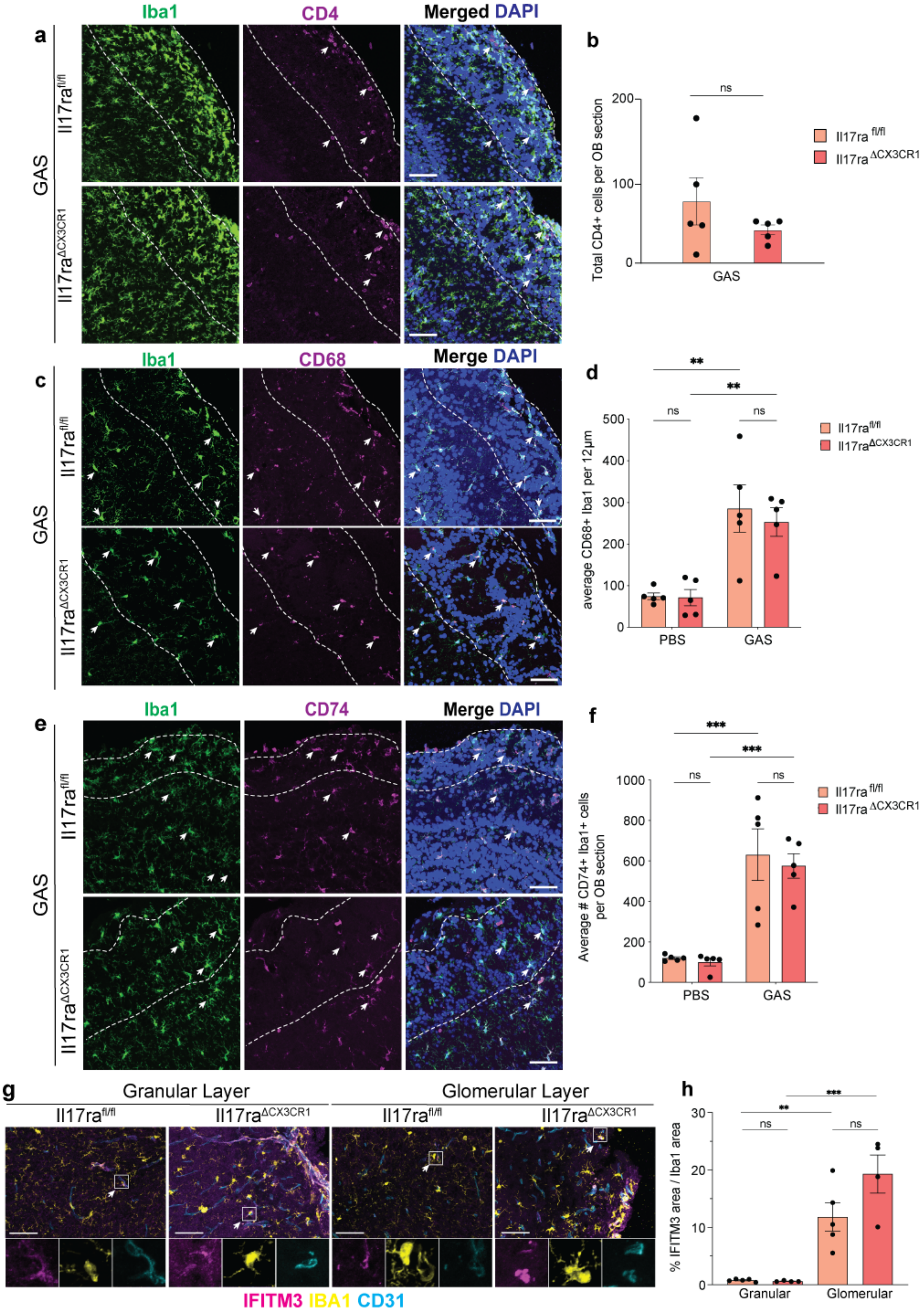

a
Iba1
CD4
Merged DAPI
GAS
Il17ra$^{fl/fl}$
Il17ra$^{\Delta CX3CR1}$
b
Total CD4+ cells per OB section
ns
GAS
Il17ra fl/fl
Il17ra ΔCX3CR1
c
Iba1
CD68
Merge DAPI
d
average CD68+ Iba1 per 12μm
PBS
GAS
e
Iba1
CD74
Merge DAPI
f
Average # CD74+ Iba1+ cells per OB section
g
Granular Layer
Glomerular Layer
IFITM3 IBA1 CD31
h
% IFITM3 area / Iba1 area
Granular
Glomerular


168

**Extended Data Figure 6. Microglia-specific deletion of the IL-17A receptor does not attenuate antigen presentation, activation, or interferon-response programs following intranasal GAS infection.**
**a, b,** Representative immunofluorescence (IF) images showing Iba1$^{+}$ myeloid cells (green) and CD4$^{+}$ T cells (magenta) in the glomerular layer of the olfactory bulb (OB; dashed outlines) from GAS-infected *Il17ra*$^{fl/fl}$ (control; salmon) and *Il17ra*$^{\Delta CX3CR1}$ (microglia/macrophage-specific knockout; red) mice, with quantification of cell numbers in the OB. Arrows indicate CD4$^{+}$ T cells in the glomerular layer. **c, d,** Representative IF images and quantification of Iba1$^{+}$CD68$^{+}$ microglia in the glomerular layer of the OB (dashed outlines) from PBS- and GAS-infected *Il17ra*$^{fl/fl}$ and *Il17ra*$^{\Delta CX3CR1}$ mice. Arrows indicate Iba1$^{+}$CD68$^{+}$ cells. **e, f,** Representative IF images and quantification of Iba1$^{+}$CD74$^{+}$ microglia in the glomerular layer of the OB (dashed outlines) from PBS- and GAS-infected *Il17ra*$^{fl/fl}$ and *Il17ra*$^{\Delta CX3CR1}$ mice. Arrows indicate Iba1$^{+}$CD74$^{+}$ cells. **g, h,** Representative IF images showing Ifitm3 (pink), Iba1 (yellow), and CD31 (blue) in the granular and glomerular layers of the OB from GAS-infected *Il17ra*$^{fl/fl}$ and *Il17ra*$^{\Delta CX3CR1}$ mice, with quantification of Ifitm3$^{+}$ area within Iba1$^{+}$ cells in the OB. Graph bars represent mean ± SEM. Statistical analysis for **b** was performed using Student's *t*-test, and analyses for **d, f, h** were performed using two-way ANOVA with Šídák's multiple-comparisons test (ns, $P > 0.05$; *$P < 0.01$; **$P < 0.001$; ***$P < 0.0001$; $n = 5$ mice per group).

**Extended Data Table 1. Batch structure of scRNA-seq experiments.** Metadata describing batch composition for all scRNA-seq samples used in this study. These samples were used to generate **Figures 1–5** and **Extended Data Figures 1–5**.

**Extended Data Table 2. Differentially expressed genes across olfactory bulb cell types in PBS- and GAS-infected wild-type mice.** Differentially expressed genes (DEGs) identified in olfactory bulb (OB) cell types comparing PBS- and GAS-infected wild-type mice. Data were used to generate **Figure 1b–d**.

**Extended Data Table 3. Differential gene expression in brain endothelial cells and microglia across experimental conditions. a,** DEGs in brain endothelial cells (BECs) and microglia from PBS- versus GAS-infected wild-type mice. Data were used to generate **Figures 1b–i, 2**, **Extended Data Figures 1a, 1c–k, 2a, 2b, 2e**, and **Extended Data Fig. 5d**. **b,** DEGs in BECs and microglia from *RORγt*$^{-/-}$ versus wild-type GAS-infected mice. Data were used to generate **Figure 4b, c, e, f** and **Extended Data Figure 4d**. **c,** DEGs in BECs and microglia from intranasally

GAS-infected $Csf2^{\Delta CD4}$ versus $Csf2^{fl/fl}$ mice. Data were used to generate **Extended Data Figure 4j, k**. **d,** DEGs in BECs and microglia from intranasally GAS-infected mice treated with anti–IL-17A monoclonal antibody versus isotype control. Data were used to generate **Figure 5b** and **Extended Data Figure 5c, d**.

**Extended Data Table 4. Gene sets used for gene set enrichment analysis. Left,** Gene sets used for GSEA in brain endothelial cells. Data were used to generate **Figure 1e**. **Right,** Gene sets used for GSEA in microglia. Dark-colored genes indicate significantly altered genes in BECs or microglia. Data were used to generate **Figures 1e and 2b**.

**Extended Data Table 5. Gene Ontology pathways enriched in brain endothelial cells and microglia. a,** GO pathways enriched in BECs and microglia from $ROR\gamma t^{-/-}$ versus wild-type GAS-infected mice. Data were used to generate **Figure 4c, f**. **b,** GO pathways enriched in BECs and microglia from $Csf2^{\Delta CD4}$ versus $Csf2^{fl/fl}$ GAS-infected mice. Data were used to generate **Extended Data Figs. 4i, 4j, 4o, 4p**. **c,** GO pathways enriched in BECs and microglia from isotype control– and anti–IL-17A monoclonal antibody–treated GAS-infected mice. Data were used to generate **Fig. 5b, c** and **Extended Data Figs. 5d, e**.

**Extended Data Table 6. Patient serum protein levels measured by multiplex immunoassay. a,** Individual patient serum cytokine and chemokine concentrations measured by multiplex immunoassay. **b,** Summary statistics for each cytokine and chemokine measured. **c,** Lower limits of detection for each analyte included in the multiplex panel. Data were used to generate **Table 1**.

**Extended Data Table 7. Patient demographic data**

Summary of patient demographic data used for serum cytokine analysis.

| | Mean age (± SD) | Sex | | Source[1] | |
|---|---|---|---|---|---|
| | | Female | Male | CUIMC | NIMH |
| **Total** | 8.7 (± 2.6) | 19 / 34 | 15 / 34 | 13 / 34 | 21 / 34 |
| **Healthy controls** | 8.7 (± 2.2) | 4 / 11 | 7 / 11 | 0 / 11 | 11 / 11 |
| **PANDAS/PANS** | 8.7 (± 2.8) | 11 / 23 | 12 / 23 | 13 / 23 | 10 / 23 |

[1] Abbreviations: Columbia University Irving Medical Center (CUIMC) and National Institute of Mental Health (NIMH)

Sample IDs and demographic information for patient cases and controls.

| Type | Sample ID[2] | Age | Sex | Source[1] |
|---|---|---|---|---|
| Healthy control | CF3 | 7.2 | Female | NIMH |
| Healthy control | CF4 | 7.6 | Female | NIMH |
| Healthy control | CF5 | 9.9 | Female | NIMH |
| Healthy control | CF7 | 12.8 | Female | NIMH |
| Healthy control | CM1 | 5.6 | Male | NIMH |
| Healthy control | CM2 | 6.8 | Male | NIMH |
| Healthy control | CM3 | 7 | Male | NIMH |
| Healthy control | CM4 | 8.3 | Male | NIMH |
| Healthy control | CM5 | 8.4 | Male | NIMH |
| Healthy control | CM6 | 10.3 | Male | NIMH |
| Healthy control | CM7 | 11.5 | Male | NIMH |
| PANDAS/PANS | PF1 | 6 | Female | CUIMC |
| PANDAS/PANS | PF4 | 8 | Female | CUIMC |
| PANDAS/PANS | PF8 | 9 | Female | CUIMC |
| PANDAS/PANS | PF9 | 9 | Female | CUIMC |
| PANDAS/PANS | PF10 | 9 | Female | CUIMC |
| PANDAS/PANS | PF2 (a,b) | 6.4 | Female | NIMH |
| PANDAS/PANS | PF3 (a,b) | 6.8 | Female | NIMH |
| PANDAS/PANS | PF5 (a,b) | 8 | Female | NIMH |
| PANDAS/PANS | PF7 (a,b) | 8.8 | Female | NIMH |
| PANDAS/PANS | PF11 (a,b) | 10.3 | Female | NIMH |
| PANDAS/PANS | PF12 | 10.6 | Female | NIMH |
| PANDAS/PANS | PM1 | 4 | Male | CUIMC |
| PANDAS/PANS | PM2 | 5 | Male | CUIMC |
| PANDAS/PANS | PM4 | 7 | Male | CUIMC |
| PANDAS/PANS | PM6 | 8 | Male | CUIMC |
| PANDAS/PANS | PM7 | 8 | Male | CUIMC |
| PANDAS/PANS | PM13 | 13 | Male | CUIMC |

[2] We obtained two separate serum samples (designated "a" and "b") for 9 of the 10 NIMH cases, which had been collected at a four- to six-week interval during their published IVIg trial (Williams et al., 2016). Overall, little difference was observed for the two timepoints in both the cytokine expression and effects of sera on HUVEC cells described in this study (**Table 1** and **Fig. 4**).

| | | | | |
|---|---|---|---|---|
| PANDAS/PANS | PM14 | 14 | Male | CUIMC |
| PANDAS/PANS | PM15 | 15 | Male | CUIMC |
| PANDAS/PANS | PM3 (a,b) | 5.7 | Male | NIMH |
| PANDAS/PANS | PM5 (a,b) | 7.4 | Male | NIMH |
| PANDAS/PANS | PM8 (a,b) | 8.9 | Male | NIMH |
| PANDAS/PANS | PM11 (a,b) | 12 | Male | NIMH |
